# Integrative multi-omics data elucidating the biosynthesis and regulatory mechanisms of furanocoumarins in *Angelica dahurica*

**DOI:** 10.1101/2024.07.23.604792

**Authors:** Jiaojiao Ji, Xiaoxu Han, Lanlan Zang, Yushan Li, Liqun Lin, Donghua Hu, Shichao Sun, Yonglin Ren, Garth Maker, Zefu Lu, Li Wang

## Abstract

Furocoumarins (FCs) are crucial natural products playing a dual role as plant defense molecules and pharmacologically active substances. *Angelica dahurica* is a renowned herb with diverse and abundant FCs. However, the accumulation pattern over developmental stages, biosynthesis pathway and regulatory mechanisms of FCs in *A. dahurica* remain elusive, hindering the production of FCs via synthetic biology approaches. Here, we constructed a chromosome-level reference genome for *A. dahurica* and quantified the content dynamics of 17 coumarins across six developmental stages of its medicinal organ, root. It showed a gradual decrease in FC concentration with root enlargement. The combined analyses of transcriptomic and metabolomic data, together with in vivo enzymatic assay, confirmed that CYP71AZ18 was involved in the biosynthesis of bergaptol, whereas CYP71AZ19 and CYP83F95 contributed to the biosynthesis of xanthotoxol. Notably, CYP71AZ19 originated from a proximal duplication event of CYP71AZ18, specific to *A. dahurica*, subsequently undergoing neofunctionalization. Accessible chromatin regions (ACRs), especially proximal ACRs, are correlated with higher gene expression levels, including the three validated genes involved in FC biosynthesis, showing potential to regulate metabolite biosynthesis. Our findings provide new insights into the biosynthetic pathway of FCs and the epigenetic regulation of metabolite biosynthesis.

## Introduction

Furocoumarins (FCs) are signific ant bioactive compounds in Apiaceae medicinal plants (Zhang *et al*., 2022). FCs played multiple roles in nature, adding a layer of complexity and charm to their research. In plants, FCs defend against pathogens, inhibit germination of competing plants, and deter herbivores (Villard *et al*., 2021). Meanwhile, FCs exhibit substantial pharmacological efficacy, encompassing anticancer, antimicrobial, and anti-inflammatory effects (Hussain *et al*., 2019; Ortega-Forte *et al*., 2021), and present promising avenues for the therapeutic management of dermatology due to its inherent photosensitizing characteristics (Jian *et al*., 2020; Richard, 2020). For instance, psoralen, bergapten and xanthotoxin plus UV-A are regarded as an invaluable therapeutic tool for psoriasis and vitiligo (Jose Serrano-Perez *et al*., 2008). Nevertheless, caution must be exercised during implementation due to the potential risk of skin allergies and interference with the metabolism of other drugs (Han *et al*., 2022).

Despite decades of study, the biosynthesis pathway of FCs remains incomplete. It involves prenyltransferases (PTs), O-methyltransferases (OMTs) and cytochrome P450s (CYP450s) (Sarker & Nahar, 2017; Rodrigues, J. L. *et al*., 2022). P450s, the most diversified enzyme family in plants (Nelson *et al*., 2004; Nelson & Werck-Reichhart, 2011), are responsible for furan-ring formation and hydroxylation. They exhibit unique characteristics, including substrate promiscuity (capacity to convert multiple substrates), catalytic promiscuity (capacity to catalyze different reactions or oxidations at different moiety of the same substrate), and multifunctionality (capacity to perform a cascade of oxidations on the same compound, which diversified into various FCs (Hansen, C. C. *et al*., 2021). A single P450 enzyme can catalyze different steps. For example, the CYP71AJ subfamily can catalyze both the conversion of marmesin to psoralen and the conversion of columbianetin to angelicin. Furthermore, FCs are primarily found in four plant families: Apiaceae, Rutaceae, Moraceae, and Fabaceae. Current research indicates that, in plants of different families, the same biosynthetic step can be catalyzed by different CYP450 families. For instance, the conversion of xanthotoxin to 5-hydroxyxanthotoxin is catalyzed by CYP71 and CYP82 families in Apiaceae, Rutaceae, and Brassicaceae, respectively. Such pattern was not observed within a single plant family. To date, CYP450s involved in the FC biosynthesis pathway belong to the CYP71, CYP82, and CYP76 families, all of which are included in the CYP71 clan (Fig. S1) (Larbat *et al*., 2009; Krieger *et al*., 2018; Jian *et al*., 2020; Villard *et al*., 2021).

On the other hand, how FC biosynthesis is regulated by epigenetic modifications is obscure. Chromatin accessibility and modification is a hallmark of regulatory DNA(Lu *et al*., 2019). Accessible chromatin regions (ACRs) reflect the gene regulatory capacity, intimately connected to gene expression patterns and metabolite accumulation (Ahmad *et al*., 2010; Yan *et al*., 2020). The chromatin accessibility landscape exhibits tissue-specific variation and undergoes dynamic changes in response to both external environmental cues and internal developmental signals (Cai *et al*., 2022). The assay for transposase-accessible chromatin with high throughput sequencing (ATAC-seq) has emerged as a powerful technology for profiling open chromatin regions across a wide range of species (Buenrostro *et al*., 2013; Buenrostro *et al*., 2015). Previous studies on chromatin accessibility in plants have primarily focused on few model plants and crops, such as *Arabidopsis thaliana* (Farmer *et al*.), rice (Wang, G *et al*., 2022), maize (Shashikant & Ettensohn, 2019), and Wheat (Pei *et al*., 2023). However, such research has been limited in medicinal plants and, in particular, how ACRs affect the accumulation of secondary metabolites is largely unexplored. Notably, the regulatory function of ACRs in artemisinin biosynthesis has been initially established in *Artemisia annua* (Zhou *et al*., 2021), a significant discovery that not only unveils the potential role of ACRs in medicinal plants but also underscores the necessity and urgency of delving deeper into their regulatory mechanisms. It is pertinent and imperative to investigate the epigenetic regulatory mechanisms governing secondary metabolite accumulation in medicinal plants through ACRs.

*Angelica dahurica* (Fisch. ex Hoffm.) Benth. & Hook. f. ex Franch. & Sav. is an essential traditional Chinese medicine (TCM) that belongs to the Apiaceae (Chinese Pharmacopoeia Commission, 2020), renowned for its medicinal and edible qualities (Huang *et al*., 2022; Wang *et al*., 2023). At least 153 FCs are reported in *A. dahurica* and FCs are the main bioactive compounds for its therapeutic efficacy, which makes *A. dahurica* an ideal system for investigating biosynthetic pathway and its epigenetic regulation of FCs (Zhao *et al*., 2022). Imperatorin and isoimperatorin are the quality control standards of *A. dahurica* in Chinese Pharmacopoeia (Chinese Pharmacopoeia Commission, 2020). Their biosynthesis precursors, bergaptol and xanthotoxol, are products derived from hydroxylation of psoralen at the C-5 (catalyzed by psoralen-5-hydroxylation, P5H) and C-8 positions (catalyzed by psoralen-8-hydroxylation, P8H), respectively. To date, only one P8H, CYP71AZ4, has been revealed in *Pastinaca sativa* (Krieger *et al*., 2018), whereas P5H remains unknown in the plant kingdom. Remarkably, P5H is the sole uncharacterized enzyme within the biosynthetic pathway leading from phenylalanine to impertinin and isoimpertinin in plants (Bourgaud *et al*., 2006; Munakata *et al*., 2021; Rodrigues, Joana L *et al*., 2022; Han *et al*., 2023). On the other hand, while *A. dahurica* holds significance in TCM, there exists a paucity of research concerning the fluctuations of FCs across its developmental stages (Liang WeiHong *et al*., 2018; Gao, H & Li, Q, 2023). This absence hindered the scientific explanation of the harvesting time for *A. dahurica*.

In this work, we present a chromosome-level genome of *A. dahurica* based on PacBio CCS and high-throughput chromosome conformation capture (Hi-C) sequencing data. We generated a comprehensive map delineating the content dynamics of 17 coumarins throughout the developmental stages of *A. dahurica* roots. By integrating transcriptomic and metabolomic datasets, our study identified and validated two P8H and one P5H involved in FC biosynthesis. Additionally, we delved into elucidating the regulatory influence of ACRs on gene expression and FC accumulation. This study completed the biosynthetic pathway from phenylalanine to imperatorin and isoimperatorin, shed light on the evolutionary history of P5H and P8H, and investigated the complex interplay between chromatin accessibility and biochemical traits.

## Materials and Methods

Additional and detailed descriptions of materials and methods were provided in Supporting Information.

### Plant materials

Seeds of *Angelica dahurica* (Fisch. ex Hoffm.) Benth. & Hook. were procured from its origin production area, Suining City (Sichuan, China), and were subsequently planted in the greenhouse of the Agricultural Genomics Institute at Shenzhen (Chinese Academy of Agricultural Sciences, China). To study the root development process of *A. dahurica*, seeds were planted in March 2021. Sampling commenced in May 2021, with a monthly interval, resulting in a total of six sampling events. Another set of *A. dahurica* plants were planted in August 2020. Upon their flowering in April 2021, five distinct tissues were sampled, including roots, stems, mature leaves, young leaves, and flowers.

*Agrobacterium tumefaciens*-mediated transient expression was performed using 4- to 5-week-old *Nicotiana benthamiana* plants. The seedlings were grown in a glasshouse under lights with a 16-hour/8-hour light/dark cycle.

### Genome assembly and annotation

The *A. dahurica* genome was assembled by integrating the sequencing data obtained with PacBio CCS and Hi-C technology using Hifiasm v0.15.5-r350 with default parameters (Table S1) (Cheng *et al*., 2021). Hi-C reads were then mapped to the contig assembly using Juicer v1.6 (Durand *et al*., 2016). A candidate chromosome-level assembly was generated automatically using the 3D-DNA v180114 (Durand *et al*., 2016). Manual check and refinement of the draft assembly was carried out via Juicebox Assembly Tools v2.18.

We used the EDTA v2.0.1 (Ou *et al*., 2019) to identify transposable elements in the *A. dahurica* genome. Coding gene structures were predicted by MAKER2 v3.01.03 pipeline (Cantarel *et al*., 2008) via three main approaches (*ab initio* predictions, homolog proteins and transcriptome data) based on the repeat-masked genome. Further annotations of protein-coding genes were conducted by GFAP (Xu *et al*., 2023) and EGGNOG-MAPPER v1.0.3 (Huerta-Cepas *et al*., 2019).

### Phylogenetic analyses

Paralogs and orthologs were identified among six Apiaceae species using the OrthoFinder v2.5.2 (Emms & Kelly, 2019), and protein sequences of single-copy orthologous genes were used to construct a phylogenetic tree. A maximum likelihood phylogenetic tree was constructed using RAxML v8.2.12 (Stamatakis, 2014). The species tree was then used as an input to estimate divergence time in the MCMCTree program in the PAML package (Yang, 2007). The expansion and contraction of gene families were inferred with CAFE5 (Mendes *et al*., 2020) based on the chronogram of the above-mentioned six plant species. To further identify the pattern of genome-wide duplications in *A. dahurica*, duplicated genes were divided using DupGen_Finder v1.12 (Qiao *et al*., 2019). Genes in the five duplicate categories were further subjected to KEGG analyses with TBtools v2.034 (Chen, C *et al*., 2023).

### Transcriptome data analyses

RNA-Seq reads of *A. dahurica* were assembled in a reference genome-based strategy with Hisat2 v2.2.1 (Kim *et al*., 2015). Gene expression was normalized as fragments per kilobase of exon model per million reads mapped (FPKM) with Stringtie v2.1.4 (Pertea *et al*., 2015). Differentially expressed genes (DEGs) were selected using DESeq2 v2_1.36.0 (Love *et al*., 2014) when |log_2_FoldChange(FC)| > 1 with adjusted *P*-value (*P*_adj_) <0.05. All genes were used for both gene ontology (GO) enrichment and Kyoto Encyclopedia of Genes and Genomes (KEGG) pathway analysis with eggNOG-mapper v2.1.9 (http://eggnog-mapper.embl.de/) (Cantalapiedra *et al*., 2021). The R package clusterProfiler v4.4.4 (Wu *et al*., 2021) was used to perform functional and pathway enrichment analysis, with *P* < 0.05 as a criterion for selecting enriched items.

### Multi-omics mining for candidate CYP450 genes

To identify potential CYP450 genes in *A. dahurica*, we integrated three distinct approaches to preliminarily obtain the CYP450 gene set. 1) We conducted a search based on conserved domains (Pfam ID: PF00067) using HMMER v3.3 (Finn *et al*., 2015); 2) we utilized a sequence similarity approach, employing the protein sequence of the CYP450 enzyme CYP71AZ4, previously characterized in *Pastinaca sativa* (Apiaceae), as a query sequence for local BLAST in the *A. dahurica* genome (--evalue 1e-6); 3) we excluded potential pseudogenes (predicted protein sequences < 300 amino acids or FPKM=0 in every sample). We considered the intersecting sequences obtained from three methods as putative CYP450 genes in *A. dahurica*. Next, we utilized a comprehensive approach that incorporated gene expression analysis, metabolite content assessment, and phylogenetic relationships to screen two candidate genes for P8H and four candidate genes for P5H.

### Functional characterization for candidate CYPs in *N. benthamiana*

Two psoralen-8-hydroxylases along with four psoralen-5-hydroxylases were cloned into the pSuper1300 vector and subsequently introduced into the GV3101 strains of *Agrobacterium* electrocompetent cells. The GV3101 strains underwent 16 h cultivation in Luria-Bertani (LB) medium supplemented with kanamycin (50 μg/mL) at 28°C and 200 rpm. Thereafter, 500 μL of the GV3101 suspension was inoculated into 30 mL of fresh LB medium containing rifampicin (25 μg/mL) and kanamycin (50 μg/mL), followed by overnight incubation. After centrifugation at 4000 rpm for 10 min, GV3101 cells were collected and resuspended in MMA buffer (comprising 100 μM acetosyringone, 10 mM MES, and 10 mM MgCl_2_, pH 5.6) for a duration of 2-3 h at ambient temperature to optical density at 600 nm (OD600) approximately 0.5-0.6. The strain was subsequently introduced into the abaxial surface of three leaves sourced from disparate 4- to 5-week-old *N. benthamiana* seedlings utilizing a needleless syringe (1 mL). After *N. benthamiana* was dark-treated for 12 h and grown under normal conditions for 3 days, psoralen at a concentration of 0.1 mM was introduced into the leaves infiltrated with *Agrobacterium* to enzymatic activity assessment.

### Identification of metabolites

Due to the photosensitivity of coumarin, all sample solutions were prepared under shaded conditions. After homogenizing the frozen tissue samples into a fine powder on a ball mill, approximately 1 g of sample powder was weighed and added to 5 mL of ethyl acetate. The mixture was vortexed for 1 min, sonicated for 30 min, and then centrifuged at 4°C and 13000 g for 15 min to collect the supernatant, which was filtered through a 0.22 μm membrane and suspended dry at -70°C in a refrigerated vacuum centrifugal concentrator. The resulting solid residue was re-solubilized with 1 mL methanol, and filtered again. 1 mL extracts were used for Liquid Chromatography-Mass Spectrometry (LC-MS) analysis using Thermo Scientific Vanquish Ultra-High Performance Liquid Chromatography System and Thermo Scientific TSQ Quantum Access Max Triple Quadrupole Mass Spectrometer (both from Thermo-Fisher Scientific)equipped with an ACQUITY UPLC®BEH C18 column (Waters Ltd., MA, USA) (2.1×100 mm, 2.5 μm).

Acetonitrile was used as mobile phase A and water with 0.1% formic acid was used as mobile phase B, and the flow rate was 0.3 mL/min. Metabolite identification was performed using authentic standards purchased from suppliers.

### Phylogenetic analysis of the CYP71 family

The construction of a phylogenetic tree incorporating both previously published and newly identified CYP genes was a pivotal step toward understanding their evolutionary relationships. Initially, experimentally verified CYP71AZ18, CYP71AZ19 and CYP83F95 genes were used as query genes to blast against *Daucus carota* L.*, Ligusticum chuanxiong* Hort. haplotype A *and Angelica sinensis* (Oliv.) Diels., *Aralia elata* (Miq.) Seem. and *Panax notoginseng* (Burkill) F. H. Chen ex C. H. Chow with E-value < 1e–5 as a cutoff. The hidden Markov model of the CYP gene (Pfam ID: PF00067) was downloaded from the Pfam database (http://pfam.xfam.org/) to further verify the identified CYP genes. CYP71A13 in *A. thaliana* were designated as outgroups. Protein sequences of these CYP genes were then aligned using MUSCLE v3.8.1551 (Edgar, 2004). Maximum likelihood trees were constructed with IQ-TREE v2.1.4 (Lam-Tung *et al*., 2015). The final phylogenetic trees were annotated and depicted with iTOL v6 (Letunic & Bork, 2016). Syntenic blocks within one species or between two species were defined by MCscanX (Wang *et al*., 2012) based on homologous gene sets with BLASTP v2.10.0.

### ATAC-seq library construction, sequencing and analysis

ATAC-seq was conducted following a previously established protocol (Lu *et al*., 2017). Briefly, 1g of frozen sample was minced in 1mL of ice lysis buffer (15 mM Tris-HCl pH 7.5, 20 mM NaCl, 80 mM KCl, 0.5 mM spermine, 5 mM 2-Mercaptoethanol, 0.2% TritonX-100). The resulting slurry containing the nuclei extract was filtered twice through a 40 μm filter. The crude nuclei containing DAPI (Sigma, D9542) were then loaded onto a flow cytometer (BD FACSCanto) for selection. After centrifugation, the nuclei pellet was washed with Tris-Mg buffer (10 mM Tris-HCl pH 8.0, 5 mM MgCl2), and Tn5 transposomes in 40 μl TTBL buffer (Vazyme, TD501) were added for a 30-minute incubation at 37°C. Following incubation, the integration products were purified using the NEB Monarch™ DNA Cleanup Kit (T1030S), and library amplification was performed using the NEB Next Ultra II Q5 master mix (M0544L). Finally, the amplified libraries were purified using Hieff NGS® DNA Selection Beads (Yeasen, 12601ES03).

Raw reads were trimmed with Fastp v0.12.4 (Chen *et al*., 2018) and then aligned to the *A. dahurica* reference genome with Bowtie2 v2.4.2 (Langmead & Salzberg, 2012). PCR duplicates, which may have arisen from amplification, were discarded with Sambamba v0.8.0 (Tarasov *et al*., 2015) and low-quality mappings (Q-valueL<L30) in the resulting BAM files were removed with SAMtools v1.13 (Li *et al*., 2009). Bigwig files were generated by deepTools v3.5.2 (Ramírez *et al*., 2016) and then imported into Integrative Genomics Viewer (IGV) v2.17.0 (Thorvaldsdóttir *et al*., 2013) to visualize the accessible chromatin landscape. ATAC peak calling was performed with MACS2 v2.2.6 (Zhang *et al*., 2008) using ‘-g 4.9e9 --keep-dup all--nomodel --shift -100 --extsize 200 -n Leaf’. CHIPSEEKER v1.16.1 (Yu *et al*., 2015) was used to retrieve the nearest genes around the peak and annotate the genomic region of the peak. The differential accessible regions (DARs) between samples were identified with MACS2 bdgdiff.

## Results

### *Angelica dahurica* genome assembly and annotation

We used PacBio sequencing in high fidelity (HiFi) with Hi-C data to assemble the genome (Table S1). Assembly of the HiFi reads yielded an initial genome of 4.89 Gb for *A. dahurica*, consistent with flow cytometry and *k*-mer analysis (Fig. S2 and S3). The assembled scaffolds are mapped onto 11 pseudochromosomes, achieving an impressive anchoring rate of 97.27% (Fig. 1; Table S2-S4). The *A. dahurica* genome size was the highest among the published Apiales assemblies (Song *et al*., 2020; Zhao *et al*., 2022; Nie *et al*., 2024). The assembly achieved 97.20%. Viridiplantae conserved genes for Benchmarking Universal Single-Copy Orthologs (BUSCO) evaluation, and LTR assembly index (LAI) was 20.81, suggesting a gold-level genome (Table. S2). Meanwhile, 81.51–96.15% of RNA-seq data were successfully mapped to the assembled genome (Table S3). All the above evidence indicates a high-fidelity genome assembly.

For genome annotation, transcriptomes of 11 tissues, including roots from seven developmental stages, stems, leaves, and flowers were sequenced (Table S2). By combining *ab initio*, homology-based and transcriptome-based approaches, we predicted 61,419 protein-coding genes (Table S5). The annotated genes exhibited high completeness with 91.00% from BUSCO, and 89.04% of them were functionally annotated through at least one of the databases NR, Pfam, GO or KEGG (Tables S5), reflecting a high level of annotation completeness. The average sequence lengths for exons, and introns are 264.69 and 606.68 bp, respectively (Table S5). In total, 4.26 Gb (87.08% of the genome) was identified as repetitive sequences, with long terminal repeat retrotransposons (LTR-RTs) being the predominant category (3.80Gb, 77.71% of the genome) (Table S6).

### Secondary metabolic pathways were enriched in duplicated genes within the expanded gene families of *Angelica dahurica*

To further investigate the genomic characteristics of *A. dahurica*, we conducted comparative genomic analysis with six other Apiaceae species, including *Centella asiatica* (L.) Urban*, Bupleurum chinense* DC*, D. carota*, *L. chuanxion* and *A. sinensie*. A phylogenetic tree of the six species based on orthologous single-copy genes revealed that *A. dahurica* diverged from *D. carota* at approximately 17.45 MYA (Fig. 1b). The phylogenetic topology is mostly consistent with previous reports (Song *et al*., 2021; Nie *et al*., 2024).

Gene family contraction and expansion analyses unveiled that 1878 and 2089 gene families contracted and expanded in the *A. dahurica* genome, respectively (Fig. 1b). The number of expanded gene families in *A. dahurica* was almost twice as that in *D. carota*, *A. sinensis* and *L. chuanxiong* (Fig. 1b). Next, genes within the expanded families in *A.dahurica* were classified into five types, whole-genome duplication (WGD), tandem duplication (TD), proximal duplication (PD), transposed duplication (TRD), and dispersed duplication (DSD) (Fig. S4). It showed that PD is the predominant type (27.89%, 1807), followed by WGD (1726, 26.64%) and TD (1190, 18.37%) (Fig. S4). All types of duplicated genes within the expanded families were enriched in primary metabolic biosynthesis (Fig. S5). Interestingly, genes classified as DSD, TD and PD within the expanded gene families were enriched in secondary metabolic pathways, such as flavonoid biosynthesis, phenylpropanoid biosynthesis and benzoxazinoid biosynthesis (Fig. S5), suggesting that dispersed, tandem and proximal duplication played vital roles in broadening the gene repertoire associated with secondary metabolic biosynthesis. Such expansions are indicative of an extensive accumulation of diverse secondary metabolites in *A. dahurica*.

### Accumulation of FCs during root development

To investigate the accumulation pattern of FCs in the root developmental stages of *A. dahurica*, we sampled *A. dahurica* roots once a month from May to October, referred to as S1 (12 weeks after seeds planting), S2 (16 weeks), S3 (20 weeks), S4 (24 weeks), S5 (28 weeks), and S6 (32 weeks), respectively (Fig. 2a). We examined root diameters and content of 17 coumarins, including 14 FCs and 3 upstream simple coumarins, which have important pharmacological effects.

The results revealed that during stages S1 to S5, the diameter of *A. dahurica* roots continued to increase and eventually stabilized at 5.2 ± 0.1 cm (Fig. 2b). As the quality control standards of *A. dahurica* in Chinese Pharmacopoeia (2020 Edition), the concentration of imperatorin and isoimperatorin remained consistently high throughout the developmental stages of *A. dahurica* (averaging 96.09 μg/g for imperatorin and 20.40 μg/g for isoimperatorin, respectively) (Fig. 2c). In addition, we found that some upstream substrates and metabolic intermediates involved in the complex FC biosynthesis exhibited distinct changes. For instance, umbelliferone, serving as the entry substance for the biosynthesis of linear and angular FCs, and marmesin, functioning as a substrate in the biosynthesis of linear FCs, showed a decreasing trend in their concentrations with root enlargement. While the concentration of xanthotoxol in *A. dahurica* roots reached the highest in the S5. In summary, despite a slight decline in the overall FC concentration with root development (Fig. 2d), the continuous increase in root diameter from S1 to S5 resulted in the highest root biomass at S5 (September), which explains the rationale for the common practice in the production of harvesting *A. dahurica* roots before bolting in the autumn season.

### Mining and verification of P8H and P5H

Multi-omics data were integrated to identify candidate CYP450 genes involved in the catalysis of psoralen to xanthotoxol and bergaptol in *A. dahurica*. Based on sequence similarity and conserved domain integrity, we identified 310 *A. dahurica* CYP450 genes, each with >300 amino acids and an FPKM > 0 in at least one tissue or root stage. Subsequently, a phylogenetic tree was constructed to compare the CYP450 genes with those from *A. thaliana* and those experimentally verified to be involved in FC biosynthesis (Bak *et al*., 2011; Krieger *et al*., 2018; Villard *et al*., 2021), resulting in 25 preliminary candidates for P8H and P5H (Fig. S6, Fig. 3c). The correlation between metabolite content and gene expression levels in five *A. dahurica* tissues (root, mature leaves, young leaves, stem, and flowers) (Fig. 3a,b) were utilized to further screen candidate genes. Ultimately, based on the correlation between FPKM and metabolite levels in diverse tissues (*r* > 0.8, Dataset S1), we selected two P8H, namely AdP8H1 (AD04G02371), AdP8H2 (AD03G00078), as well as four P5H: AdP5H1 (AD04G02366), AdP5H2 (AD08G00823), AdP5H3 (AD03G00273), and AdP5H4 (AD04G04017) (Fig. 3c) for downstream experimental validation. Among these genes, AdP8H1, AdP8H2, and AdP5H1 exhibited corresponding catalytic activities in a tobacco transient expression system (Fig. 3d), named them CYP71AZ19, CYP83F95, CYP71AZ18, respectively.

### CYP71AZ19 results from a *A.dahurica*-specific proximal duplication of CYP71AZs

To delve into the evolutionary history of CYP71AZs in *A.dahurica*, we collected three groups of CYP71 genes: 1) three verified *A.dahurica* genes, CYP83F95, CYP71AZ18 and CYP71AZ19; 2) CYP71s from three Apiaceae species (*D. carota*, *L. chuanxiong* and *A. sinensis*) and two Araliaceae species (*A. elata* and *P. notoginseng*) queried by three aforementioned *A.dahurica* genes; (3) CYP71As and CYP71Bs in *A. thaliana* representing outgroup. The phylogenetic tree of CYP71s was divided into two clades (Fig. 4a). Clade I represented the CYP71AZ subfamily within the Apiaceae and Araliaceae families, including CYP71AZ18 and CYP71AZ19; while Clade II encompassed the CYP83F subfamily in these families, featuring CYP83F95 (Fig. 4a). The topological structures of Clade I and II were strikingly similar, with Araliaceae CYP residing on the basal branches, and more copies of CYP71AZs and CYP83Fs were observed in Apiaceae than in Araliaceae (Fig. 4a). It suggests that both CYP71AZ and CYP83F subfamily underwent lineage-specific duplication events in Apiaceae post its divergence with Araliaceae.

Based on the synteny analysis, CYP71AZ18 was collinear with AS02G00065 in *A. sinensis*, DCAR003921 in *D. carota*, LCX8AG000448 in *L. chuanxiong*, respectively (Fig. 4b). However, no such collinearity was observed for CYP71AZ19 with any genes in the above-mentioned Apiaceae species (Fig. 4b). Coupled with the finding that CYP71AZ18 and CYP71AZ19 are products of proximal duplication (Fig. S3), it can be inferred that CYP71AZ19 originated from a proximal duplication event of CYP71AZ18 during the speciation of *A. dahurica*.

### The landscape of chromatin accessibility in *Angelica dahurica*

ATAC-Seq was applied to investigate the chromatin accessibility of different tissues and during the root developmental stages of *A. dahurica*. By plotting the positions of chromatin accessibility relative to all genes (including coding regions and 3 kb upstream and downstream regions), we found that chromatin accessibility was enriched in transcription start sites (TSS) and transcription end sites (TES) (Fig. 5a,b, Fig. S7a,b), showing a similar distribution pattern in leaves and roots at different developmental stages. We next identified 3833, 33 211, 29 410, 42 281, and 56 543 ACRs in leaves and roots (S0, S1, S3 and S5), respectively. These were assigned to the nearest genes based on the annotation. Among 10 genomic elements (promoter (≤ 1 kb), promoter (1-2 kb), promoter (2-3 kb), 3’UTR (3’un-translated region), 1st exon (first exon), other exons, 1st intron (first intron), other introns, downstream, distal intergenic regions, 5’UTR), the distal intergenic regions were the most annotated type, which were more abundant in roots (58.03%-79.69%) than in leaves (49.84%), followed by the promoter region (≤1 kb) (Fig. 5c). Considering that each genomic element has different proportions on the genome, we further divided the genome into 6 parts, TSS (TSS±1 kb), TES (TES±1 kb), gene, exon, intron, and intergenic regions, for further enrichment analyses of ACRs, and found that it was mainly enriched in exon, TSS and TES regions, whereas intergenic showed negative enrichment. (Fig. 5d, Fig S8).

We further noticed that genes with ACRs had higher expression levels than genes without ACRs in both leaves and roots (*P* < 0.0001) (Fig. 5e, Fig S9). To explore the effects of ACRs in different regions of the genome on gene expression, we categorized ACRs into proximal ACRs (located in 3 kb upstream and 300 bp downstream of the gene) and distal ACRs (located in intergenic regions: > 3 kb upstream of the gene or > 300 bp downstream of the gene). The results showed that genes with both proximal and distal ACRs exhibited the highest expression levels, followed by those possessing only proximal ACRs. Moreover, all three categories of ACR-associated genes showed significantly higher expression levels compared to genes lacking ACRs (*P* < 0.0001) (Figure 5f).

In addition, we identified differential accessible regions (DARs) between roots and young leaves, with 6494 DARs up-regulated in roots, significantly enriched in the phenylpropanoid biosynthesis pathway (ko00940), and 2381 DARs up-regulated in leaves, significantly enriched in photosynthesis (ko00195) (Fig. S10). According to gene expression data, 5620 and 6331 differentially expressed genes (DEGs) were up-regulated in roots and young leaves, respectively. Notably, among these, 1760 root highly expressed genes overlapped with root specific ACRs, accounting for 27.10% of the root specific ACRs, and were notably enriched in secondary metabolic pathways, such as phenylpropanoid biosynthesis pathway (ko00940), pertinent to coumarin biosynthesis (Fig. 5g,h). In contrast, only 295 leaf highly expressed genes overlapped with leaf specific ACRs, accounting for 12.39% of the leaf specific ACRs and significantly enriched in the photosynthesis (ko00195) (Fig. 5g,h). Taken together, ACRs are crucial in regulating gene expressions, especially in the phenylpropanoid biosynthesis pathway in roots (upstream pathway of coumarin biosynthesis).

### Chromatin accessibility is associated with FC biosynthesis regulation in *Angelica dahurica*

To illustrate the possible regulation of chromatin accessibility on the expression of genes related to FC biosynthesis, we further focused on the relationship between gene expression and chromatin accessibility in roots and leaves for the three validated genes. Compared with leaves, CYP71AZ19 and CYP71AZ18 showed higher chromatin accessibility and gene expression in roots, whereas CYP83F95 had higher chromatin accessibility in roots although its expression level was similar in roots and leaves (Fig. 7a). In addition, genes with high chromatin accessibility in roots enriched some TF binding motifs associated with stress response, such as abscisic acid-insensitive 3 (ABI3) facilitating plant adaptation to abiotic stressors, as well as WRKY aiding in plant defense against pathogen (Fig. 7b), which is consistent with the biodefense function of FCs.

Focusing on the broader FC biosynthetic pathway, we employed genes with experimentally validated catalytic functions as query (Dataset S2). Subsequently, genes exhibiting over 75% homology to queries were identified as candidate genes, and their expression levels and chromatin accessibility were quantified. Our findings revealed that while the expression patterns of these genes in roots and leaves did not consistently mirror the trends in metabolite content, they consistently correlated with chromatin accessibility. Notably, across nearly every step of the pathway, genes exhibiting consistent chromatin accessibility, expression levels and downstream product concentrations were observed. These genes represent promising candidates for subsequent experimental validation and merit further investigation as primary targets.

## Discussion

Here, we constructed a high-quality chromosome-level reference genome of *A. dahurica.* Bioinformatic analysis, along with in vivo enzymatic assay, confirmed that CYP71AZ18 is involved in the biosynthesis of bergaptol, whereas CYP71AZ19 and CYP83F95 contribute to the biosynthesis of xanthotoxol, elucidated the biosynthetic pathway leading from phenylalanine to imperatorin and isoimperatorin. Notably, CYP71AZ19 originated from a proximal duplication event of CYP71AZ18 on the speciation of *A. dahurica*, subsequently undergoing neofunctionalization. The ATAC-seq analysis demonstrated that chromatin accessibility was positively correlated with gene expression, especially in the proximal ACRs. Notably, the expression of FC biosynthesis genes was also regulated by chromatin accessibility. Furthermore, we investigated the accumulation patterns of 17 coumarins during the developmental stages of *A. dahurica* roots, and found a gradual decrease in FC concentration as the roots developed. Our findings provide new insights into the biosynthetic pathway of FCs, its evolutionary history, and the epigenetic regulation of secondary metabolite biosynthesis.

Understanding the accumulation pattern of important pharmacologically active compounds in medicinal plants throughout their developmental stages is crucial for optimizing the harvest time and enhancing resource utilization(Wang, Q *et al*., 2022). Our study reveals that the concentration of imperatorin and the 17 coumarins peaked at the S1 period (12 weeks), whereas isoimperatorin reached its maximum level at the S2 period (16 weeks), which is partially consistent with Liang et al.’s observation that the highest concentrations of imperatorin and isoimperatorin peaked at 23 weeks of root growth instead of the harvest time in *A. dahurica* (Liang WeiHong *et al*., 2018). This decline may be attributed to the predominant distribution of coumarins in the phloem of the roots (Gao, H & Li, QJPA, 2023), which becomes relatively contracted as the proportion of xylem increases with root enlargement, resulting in a decrease in the concentration of coumarins per unit volume. However, given that the diameter of *A. dahurica* roots continued to increase during the S1-S5 period, with a corresponding increase in root biomass, harvesting during the S5 period remains sensible. Moreover, due to biosynthesis of secondary metabolites demands significant energy expenditure and plants tend to balance energy consumption between growth and defense (Alba *et al*., 2012; Hunziker *et al*., 2021), the observed high content of FCs at the early developmental stage of roots might indicate higher significance of biodefense at the seedling stage than that at mature stages. The sustained high levels of imperatorin and isoimperatorin observed throughout root development may reflect their pivotal role in biodefense.

The CYP450s constitute an old enzyme superfamily that can be found in nearly all organisms (Cresnar & Petric, 2011). They typically catalyze monooxygenation/oxidation reactions, and their diversification plays a crucial role in enabling the construction of numerous complex molecules and is a key driving force of phytochemical diversity (Mizutani & Sato, 2011; Hansen, Cecilie Cetti *et al*., 2021). In the FC biosynthesis of *A. dahurica*, three genes from the CYP71 clan, CYP71AZ18, CYP71AZ19 and CYP83F95 respectively react on hydroxylation at C-5 and C-8 positions of psoralen, significantly contribute to the final diversity of FCs in *A. dahurica* (Fig. 4). The identification of CYP71AZ18 completes bergaptol biosynthesis pathway, a gap in FC biosynthesis pathway. In addition, the functional convergence observed between CYP71AZ19 and CYP83F95, despite their classification into distinct families, underscores the importance of broadening the thresholds in candidate gene screening based on phylogenetic relationships. Several reports highlighted that lineage-specific tandem duplications and diversifications frequently occurred in the CYP71 clan, elucidating that the genetic redundancy ensuing from duplication events serves as a reservoir for functional innovation (Schenck & Last, 2020; Liu *et al*., 2021). Our results revealed a pattern of *A. dahurica*-specific proximal duplication for driving the quick formation and diversification of CYP71AZ members (Fig. 5). Such expansions enrich the FC content and phytochemical diversity in *A. dahurica*, which are commonly assumed to be associated with adapting to local environments or defending biotic stress (Hamberger & Bak, 2013). To sum it up, the identification and evolutionary history of CYP71AZ provides an exciting and novel example of how P450 family expands and evolves, leading to its recruitment in rich content and abundant diversity of FCs.

Chromatin accessibility is crucial for maintaining precise regulation of gene expression (Buenrostro *et al*., 2013; Buenrostro *et al*., 2015; Guo *et al*., 2021; Chen, G *et al*., 2023). The regulatory elements controlling gene expression can be broadly categorized into three types: promoters, enhancers, and boundary elements (such as insulators and boundary enhancer), among which, promoters are primarily located at the TSS while enhancers and boundary elements are often situated at distal positions away from genes (Oudelaar & Higgs, 2021). The distribution of ACRs in *A. dahurica* is notably enriched in the proximal region of genes, mainly exon, TSS and TES. And these proximal ACRs have a stronger impact on gene expression compared with those distal counterparts, which implies that a potentially dominant role of proximal ACRs in transcriptional regulation through the regulation of promoter sequences and transcriptional accessibility of gene regions, which aligns with previous research (Lu *et al*., 2019; Zhou *et al*., 2022). Furthermore, genes with both proximal and distal ACRs exhibited the highest expression levels imply a synergistic regulatory interplay between proximal and distal ACRs in modulating gene expression. Moreover, metabolite-enriched roots exhibited a larger number of DARs and DEGs with consistent alteration patterns compared to leaves. Genes associated with up-regulated DARs and up-regulated expression levels in roots were enriched in diverse metabolic pathways, including phenylalanine biosynthesis and phenylalanine metabolism (upstream pathway of coumarin biosynthesis). The three experimentally verified CYP450s displayed a higher chromatin accessibility pattern in roots than leaves, concomitant with the distribution trends of catalytic products, which further corroborates the pivotal role of chromatin accessibility in regulating the synthesis of secondary metabolites in plants. This corroborates the significant influence of chromatin accessibility on secondary metabolite synthesis, as well as plant growth and development. Given that biotic and abiotic stresses can induce the biosynthesis of FCs in *A. dahurica*, stress-responding related motifs were enriched among the root up-regulated ACRs. Our study establishes a theoretical foundation for leveraging epigenetic regulation to enhance the biosynthesis of FCs.

In conclusion, our study provides a valuable genomic resource of *A. dahurica*, enriches the secondary metabolite profile during its root development, fills a gap in the FC biosynthetic pathway, explores the evolution of the FC biosynthetic genes, and dissects the epigenetic regulation of gene expression and metabolite biosynthesis. This research advances our knowledge of the FC biosynthesis pathway and the role of lineage-specific duplication of CYP450 in diversifying FCs, and contributes to the understanding of the impact of epigenetic regulation on gene expression, which provides insights into metabolic production of FCs via biosynthetic technology.

## Acknowledgements

We thank the support from Innovation Program of Chinese Academy of Agricultural Sciences. We also thank D. Nelson of P450 nomenclature committee for naming the CYPs. This work was supported by the National Key Research and Development Program of China, grant 2023YFA0915800; National Natural Science Foundation of China, grants 32300223, 32070242, and 82373837; Shenzhen Fundamental Research Program, grant 20220817165436004; Shenzhen Science and Technology Program, grant KQTD2016113010482651; Key Project at Central Government Level (The ability establishment of sustainable use for valuable Chinese medicine resources), grant 2060302; Special Funds for Science Technology Innovation and Industrial Development of Shenzhen Dapeng New District, grants RC201901-05 and PT201901-19; China Postdoctoral Science Foundation, grant 2020 M672904; Basic and Applied Basic Research Fund of Guangdong, grant 2020A1515110912; Science, Technology, and Innovation Commission of Shenzhen Municipality of China, grant ZDSYS20200811142605017.

## Author contributions

LW conceived and designed the study. JJ prepared the materials. XH assembled and annotated the genome, and conducted WGD analysis. YL and ZL performed the ATAC experiment. JJ performed ATAC analysis, screened candidate genes and visualization. ZL and DH cloned and characterized the candidate genes. JJ, LL and XH wrote the manuscript. LW, ZL, SS, YR and GM revised the manuscript. All authors read and approved the final manuscript. JJ and XH contributed equally to this work.

## Data availability

All data needed to evaluate the conclusions in the paper are present in the paper and/or the Supplementary Materials. The data that support the findings of this study, including the raw genome, RNAseq and ATAC-seq data, were deposited in the CNGB Nucleotide Sequence Archive (https://db.cngb.org/cnsa) and are accessible with the accession ID CNP0005587.

## ORCID

Jiaojiao Ji: https://orcid.org/0009-0006-5974-7389

Xiaoxu Han: https://orcid.org/0009-0007-2542-6402

Lanlan Zang:

Yushan Li:

Liqun Lin: https://orcid.org/0009-0007-5490-9091

Donghua Hu:

Shichao Sun:

Zefu Lu: https://orcid.org/0000-0001-9322-8351

Yonglin Ren: https://orcid.org/0000-0003-4091-8812

Garth Maker: https://orcid.org/0000-0003-1666-9377

Li Wang: https://orcid.org/0000-0003-2068-7535

---

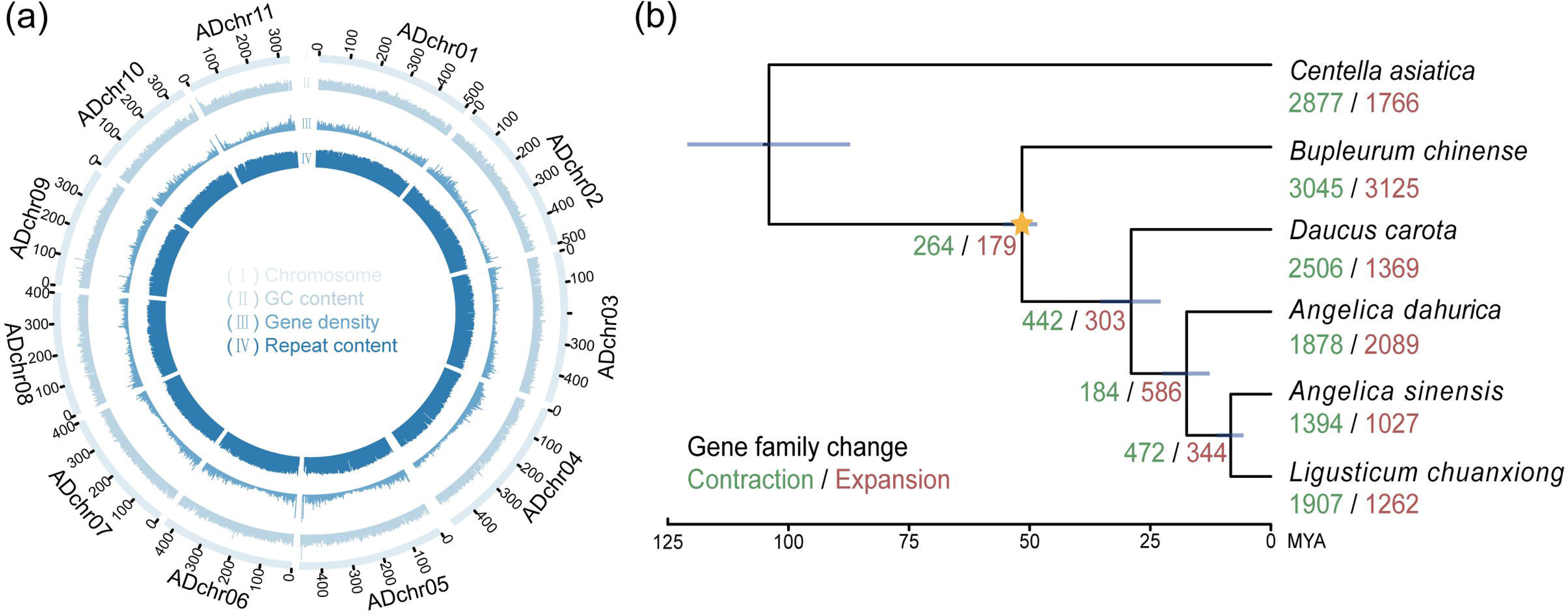

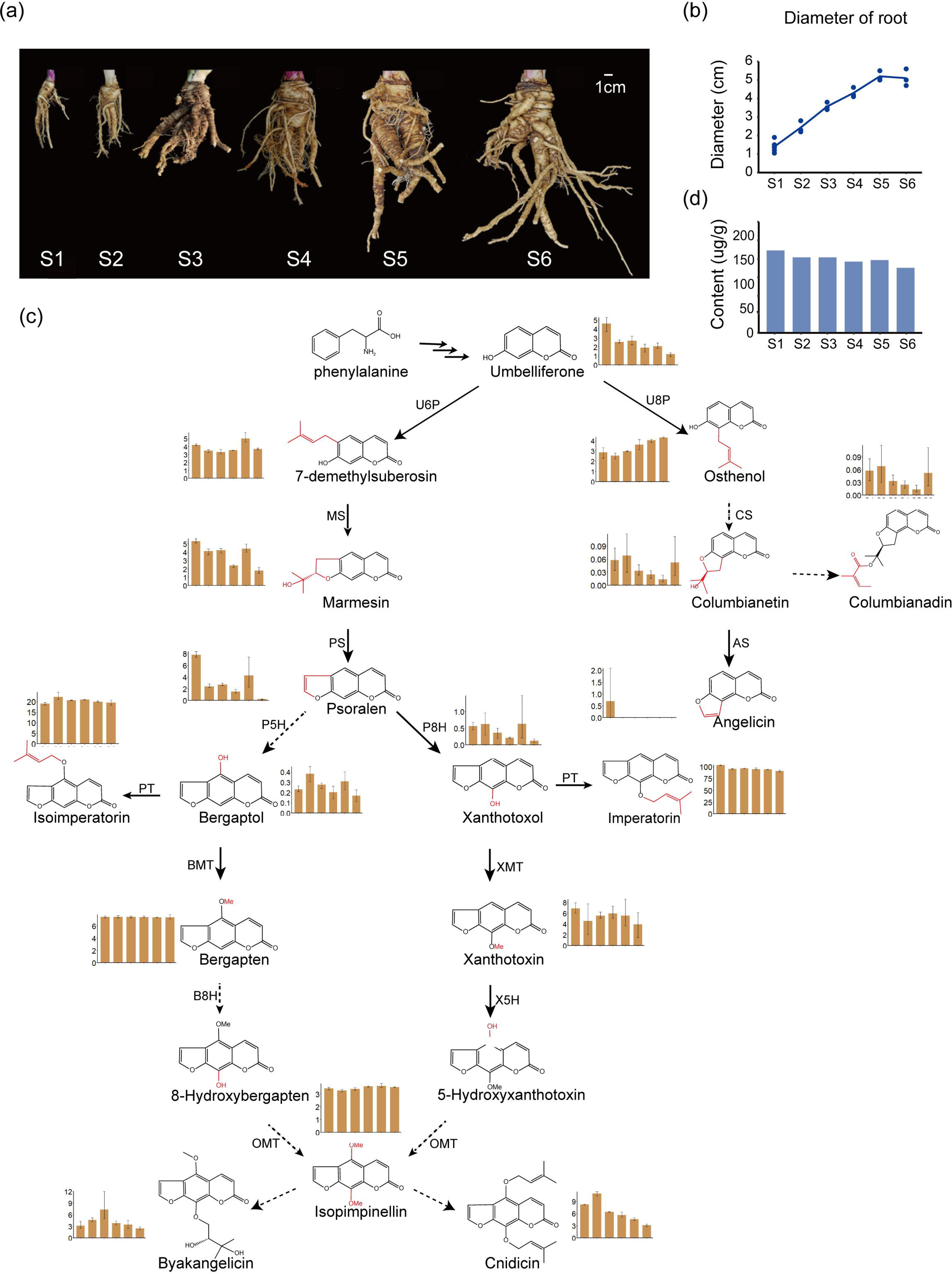

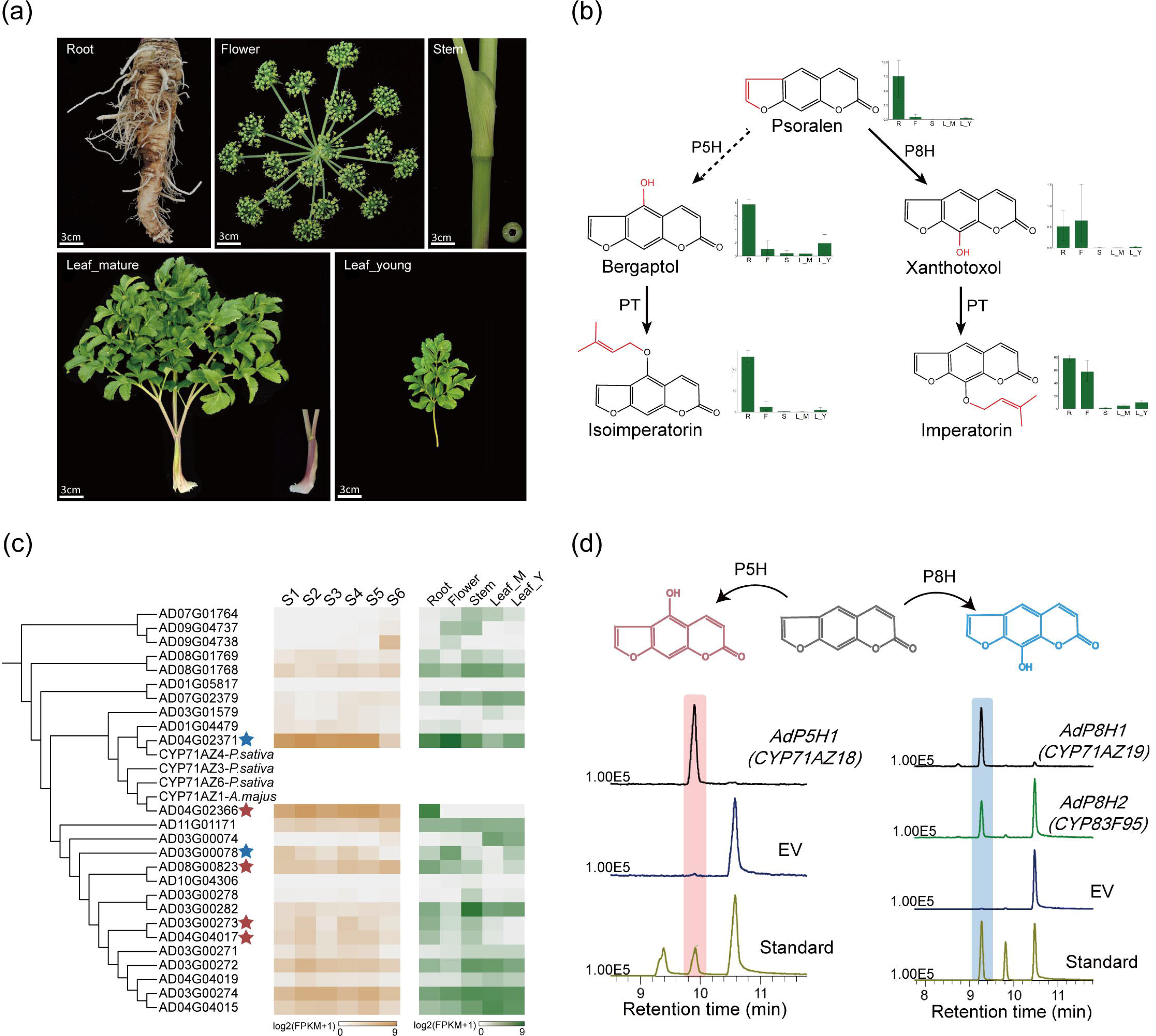

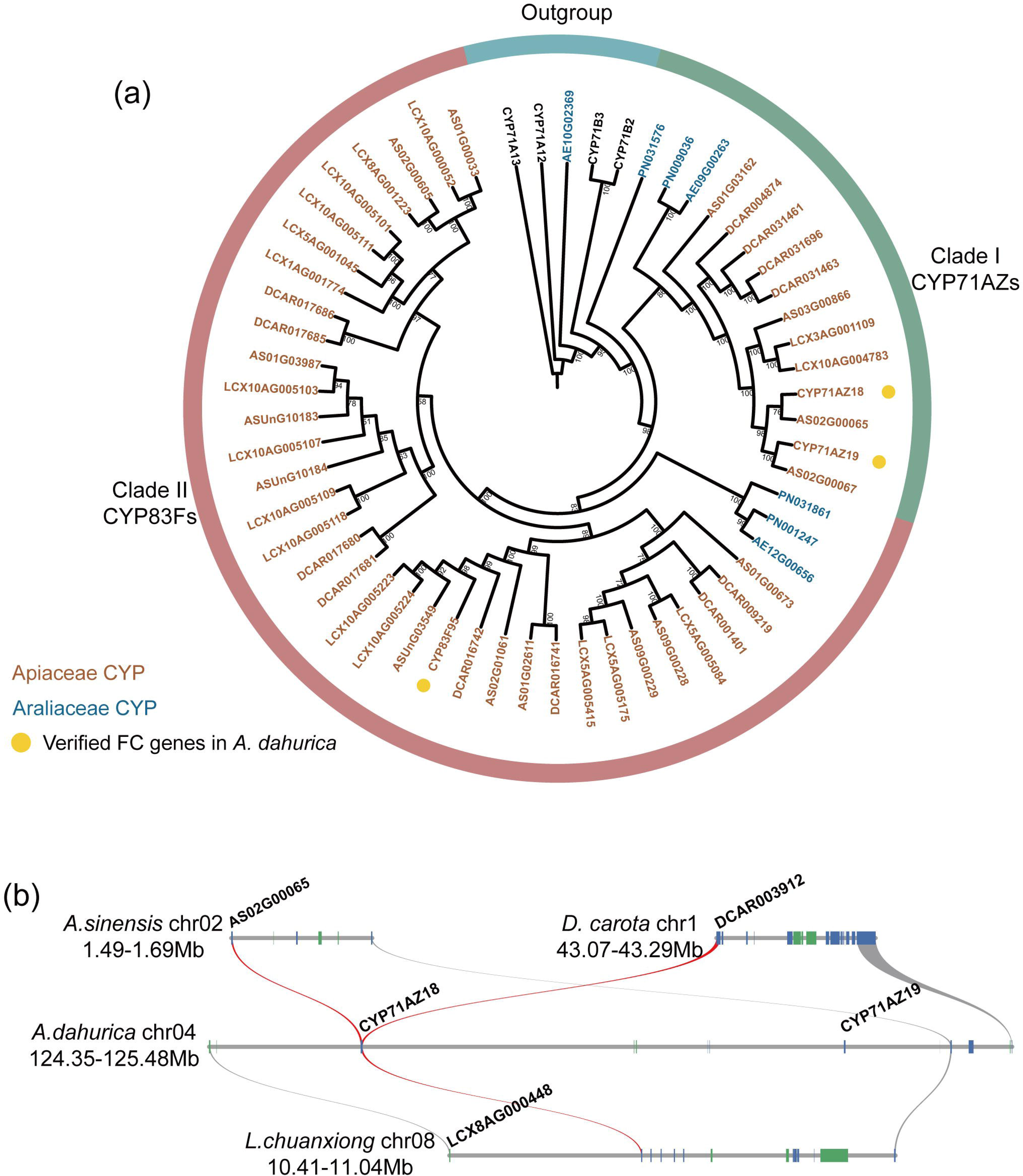

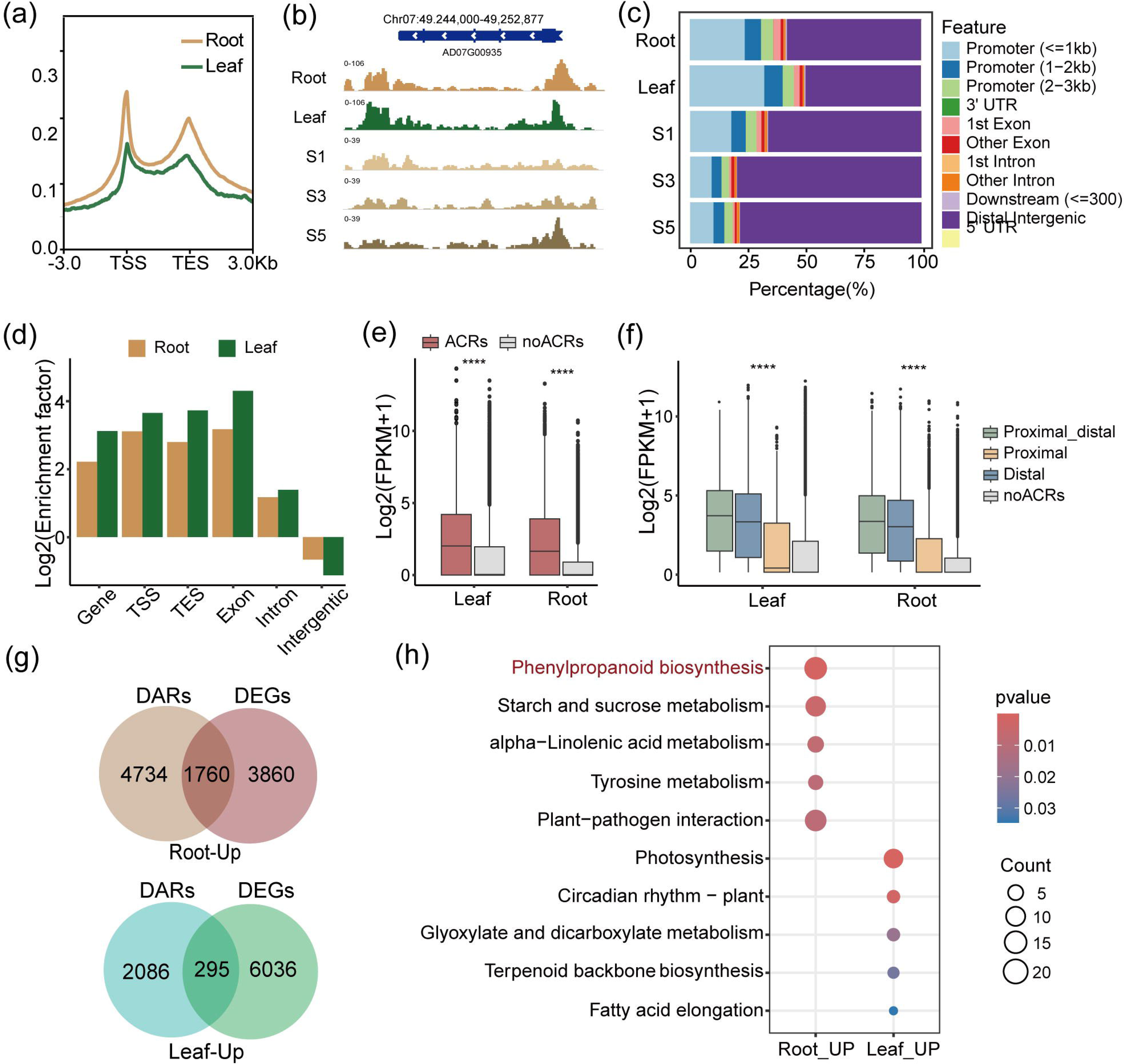

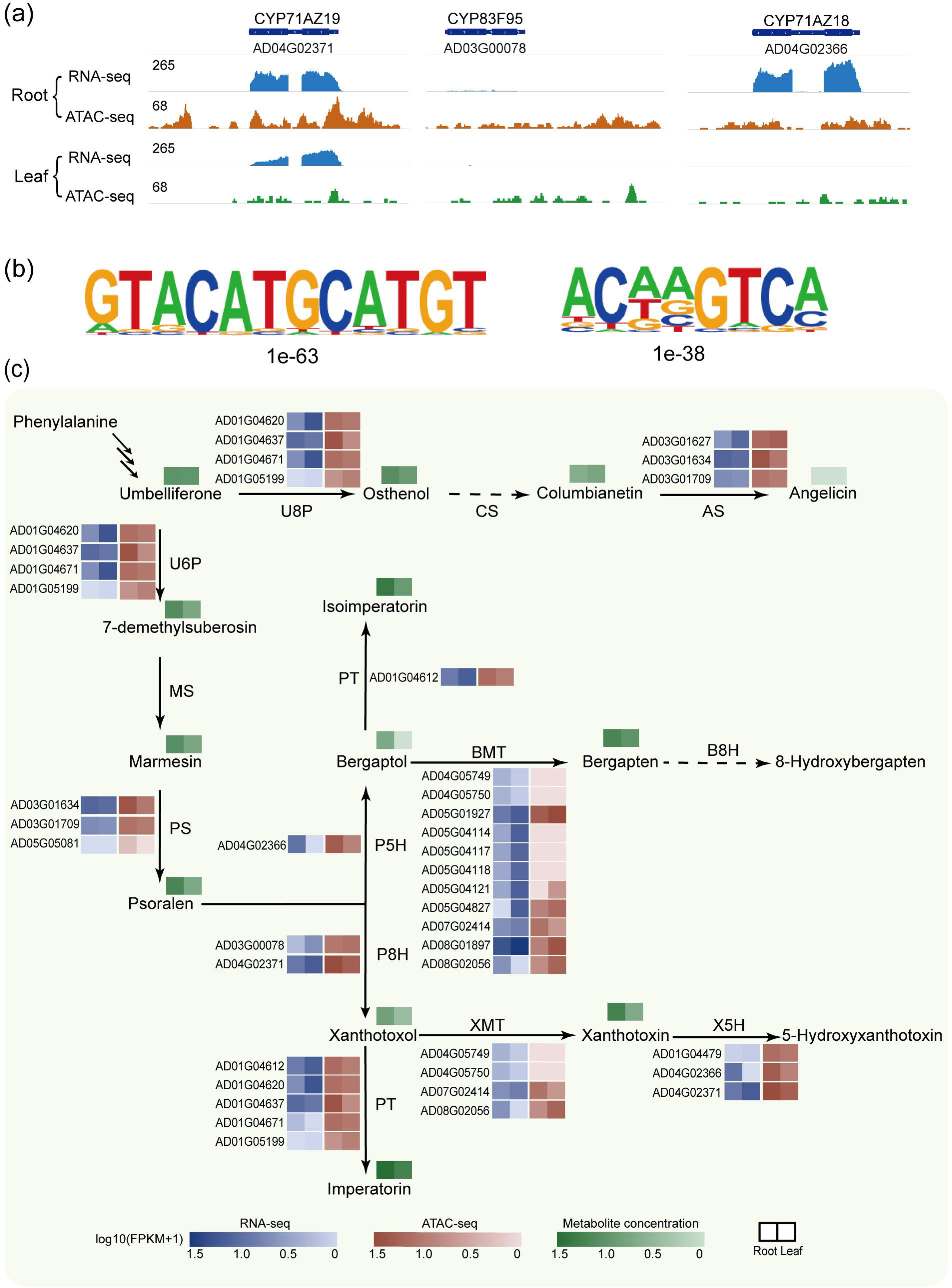

