## Supplemental Figure, Table and Methods for "Integrative multi-omics data elucidating the biosynthesis and regulatory mechanisms of furanocoumarins in *Angelica dahurica*"

***New Phytologist* Supporting Information**

**Article acceptance date:**

The following Supporting Information is available for this article:

**Dataset S1** Pearson correlation between FPKM and metabolite levels in diverse tissues.

**Dataset S2** Validated genes in the FC biosynthetic pathway.

**Fig. S1** Schematic representation of the furanocoumarin biosynthetic pathway and the enzymatic steps catalyzed by cytochrome P450s.

**Fig. S2** The K-mer depth distribution for genome size evaluation.

**Fig. S3** Flow cytometry with *A. sinensis* and *Foeniculum vulgare* as reference.

**Fig. S4** Classification of duplicated genes within expanded gene families in *A. dahurica*.

**Fig. S5** Kyoto Encyclopedia of Genes and Genomes (KEGG) enrichment analyses of different types of duplicated genes within expanded gene families.

**Fig. S6** Phylogenetic tree of CYPs from *A. dahurica* for screening candidate genes responsible for FC biosynthetic pathway.

**Fig. S7** Chromatin accessibility profiles in gene regions.

**Fig. S8** Bar plot of enrichment factor of accessible chromatin regions (ACRs) in gene, TSS (within 1 kb upstream and downstream of the TSS), TES (within 1 kb upstream and downstream of the TES), exon, intron, and intergenic regions.

**Fig. S9** Expression levels of genes with or without related ACRs in root development.

**Fig. S10** KEGG enrichment analysis of differential accessible regions (DARs) up-regulated in leaves (left) and root (right) of *A.dahurica*.

**Table S1** Sequences generated in this study.

**Table S2** Assembly and annotation statistics of the Angelica dahurica genome.

**Table S3** The mapping rates of A. dahurica in different tissues.

**Table S4** Statistics of Angelica dahurica pseudomolecules.

**Table S5** Annotation statistics of the Angelica dahurica genome.

**Table S6** Repeat contents in the Angelica dahurica genome.

**Methods S1** Genome size estimation.

**Methods S2** DNA and RNA preparation and sequencing.

**Methods S3** Genome assembly and annotation.

**Methods S4** Phylogenetic analyses.

**Methods S5** Multi-omics mining for candidate CYP450 genes.

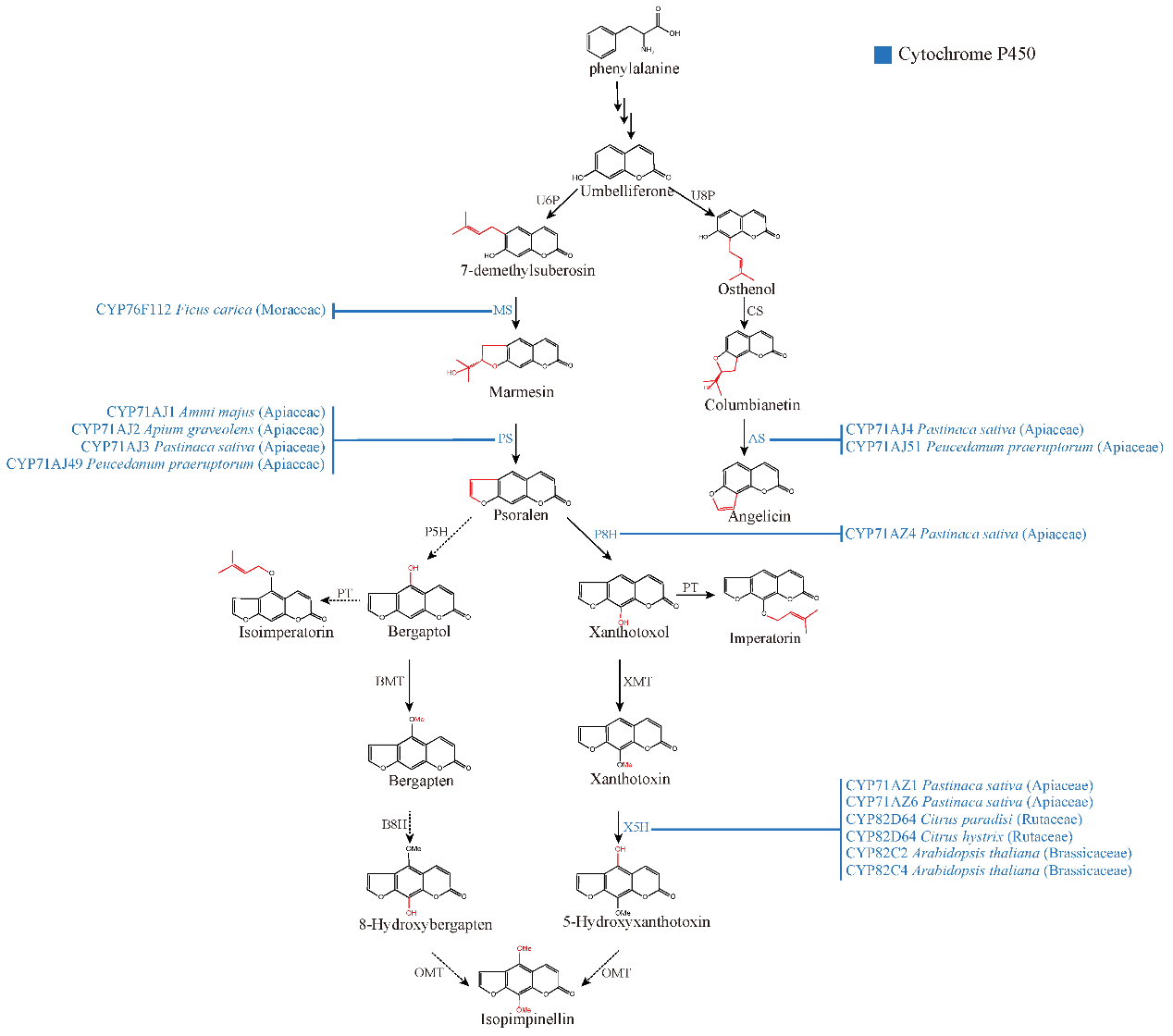

**Fig. S1** Schematic representation of the furanocoumarin biosynthetic pathway and the enzymatic steps catalyzed by cytochrome P450s.

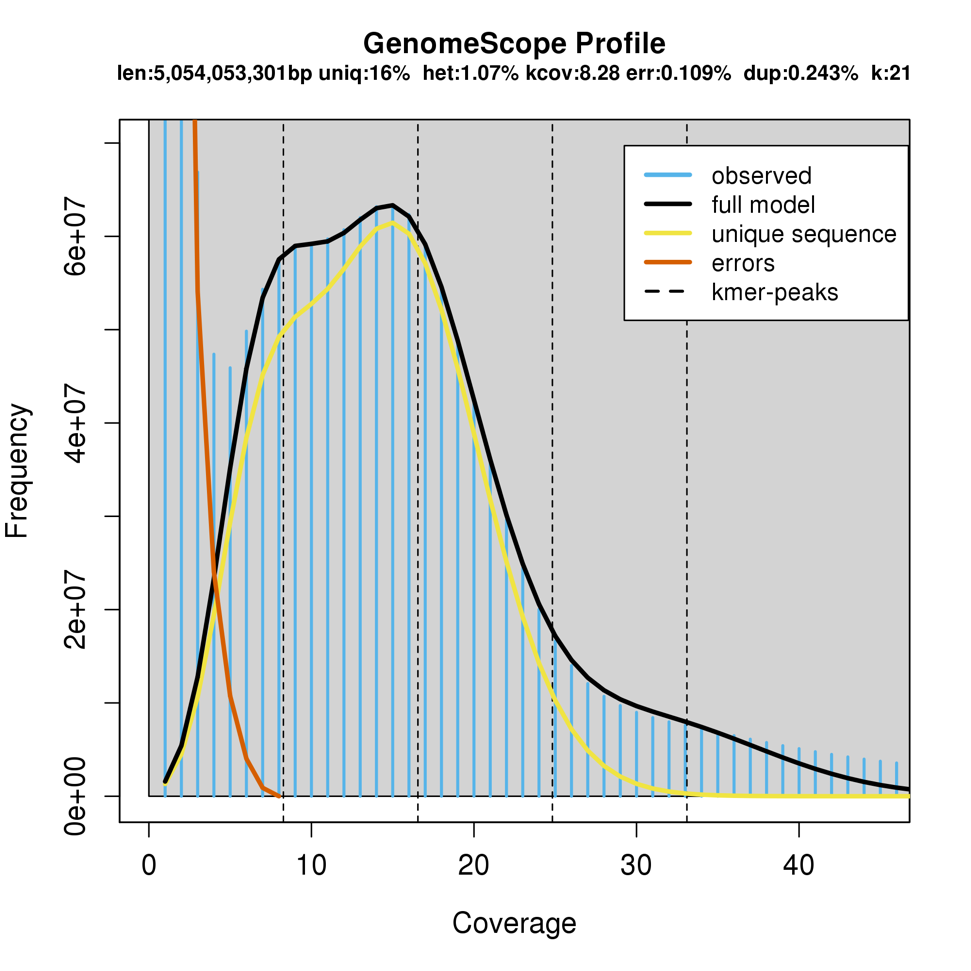

**Fig. S2** The K-mer depth distribution for genome size evaluation.

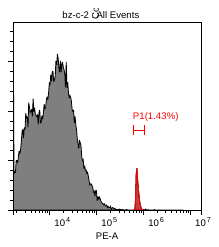

**Fig. S3** Flow cytometry with *A. sinensis* and *Foeniculum vulgare* as reference. Genome size was calculated as about 4.56 Gb.

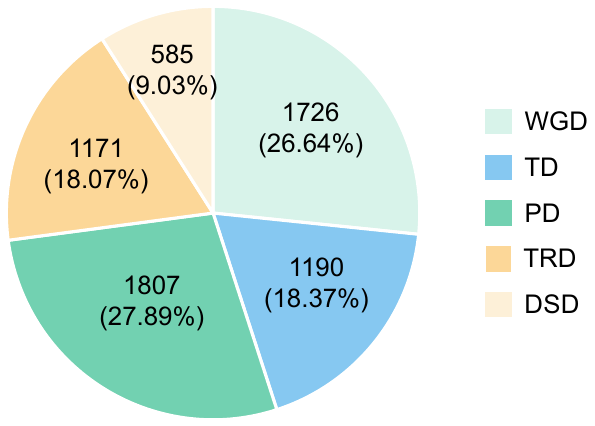

**Fig S4** Classification of duplicated genes within expanded gene families in *A. dahurica*. WGD, whole-genome duplication, TD, tandem duplication, PD, proximal duplication, TRD, transposed duplication, DSD, dispersed duplication.

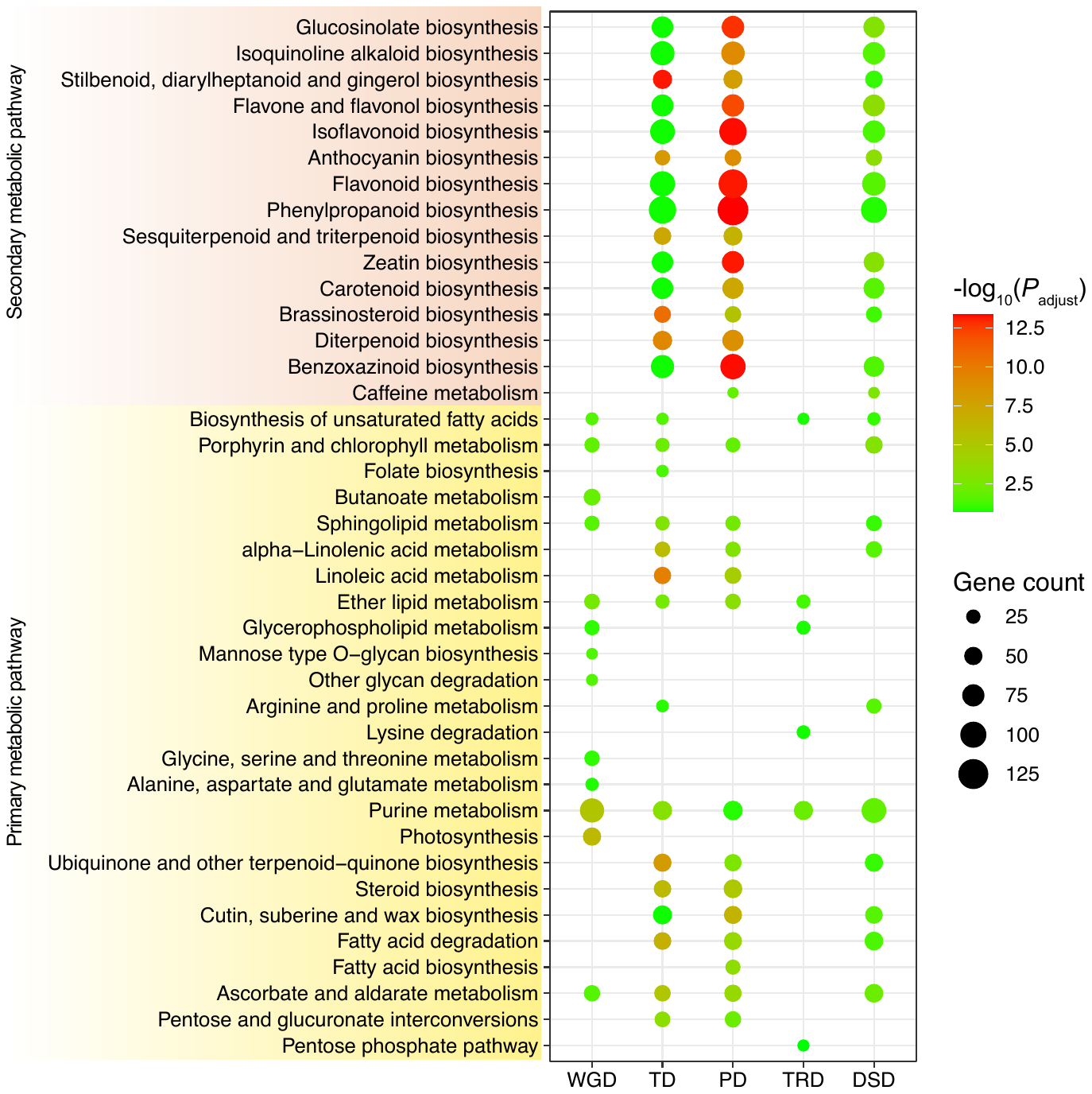

**Fig. S5** Kyoto Encyclopedia of Genes and Genomes (KEGG) enrichment analyses of different types of duplicated genes within expanded gene families. The enriched terms with adjusted *P* < 0.05 are presented. Color of the bubbles indicates statistical significance of the enriched terms; size of the bubbles indicates number of genes within the term.

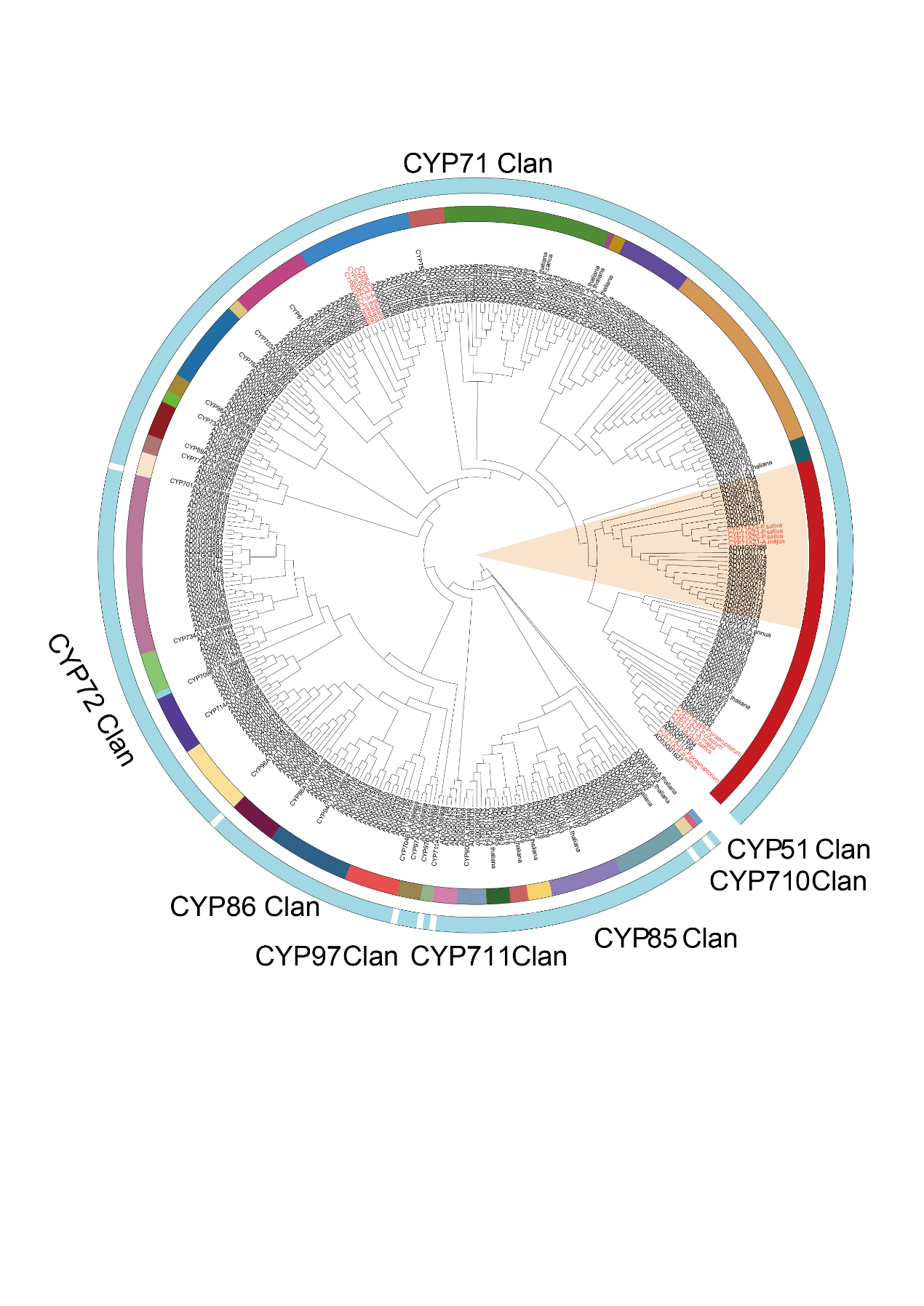

**Fig. S6** Phylogenetic tree of CYPs from *A. dahurica* for screening candidate genes responsible for FC biosynthetic pathway. The orange shaded lineages represent 25 preliminary candidate genes for P8H and P5H. The red font represents experimentally validated genes involved in furanocoumarin biosynthesis.

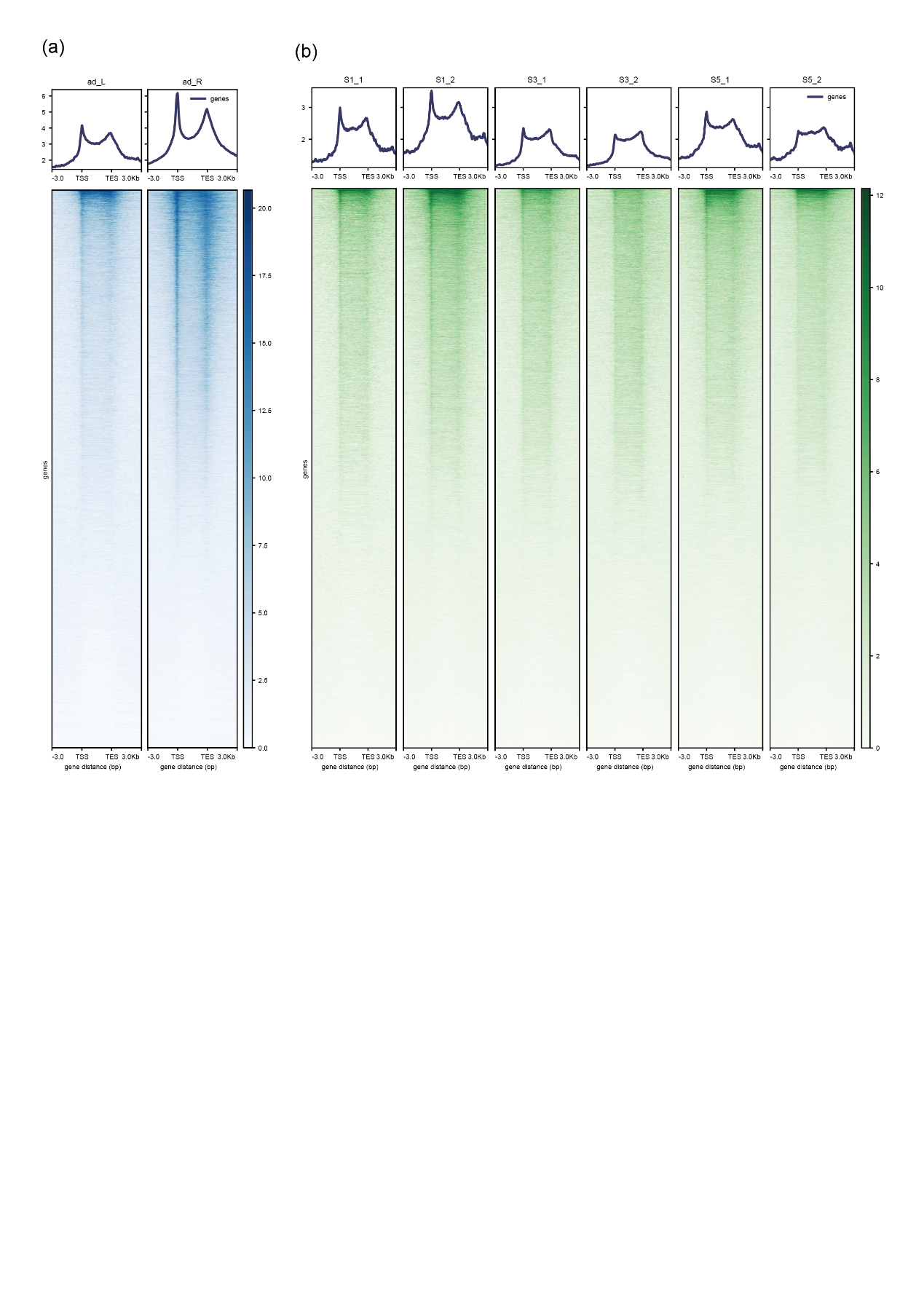

**Fig. S7** Chromatin accessibility profiles in gene regions. The 3 kb upstream and downstream flanking coding regions were aligned for all genes. TSS, transcription start site; TES, transcription end site.

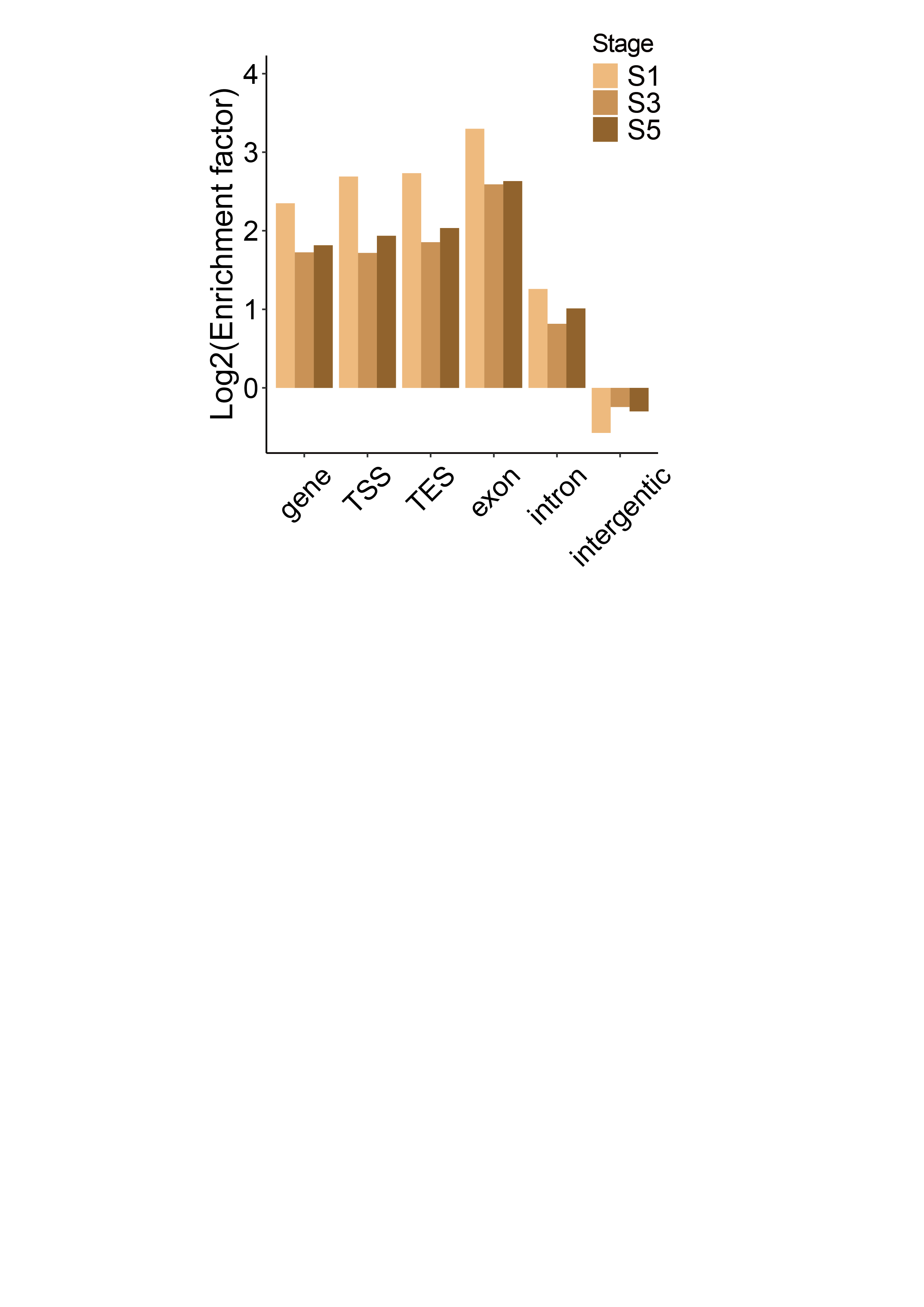

**Fig. S8** Bar plot of enrichment factor of peaks (ACRs) in gene, TSS (within 1 kb upstream and downstream of the TSS), TES (within 1 kb upstream and downstream of the TES), exon, intron, and intergenic regions.

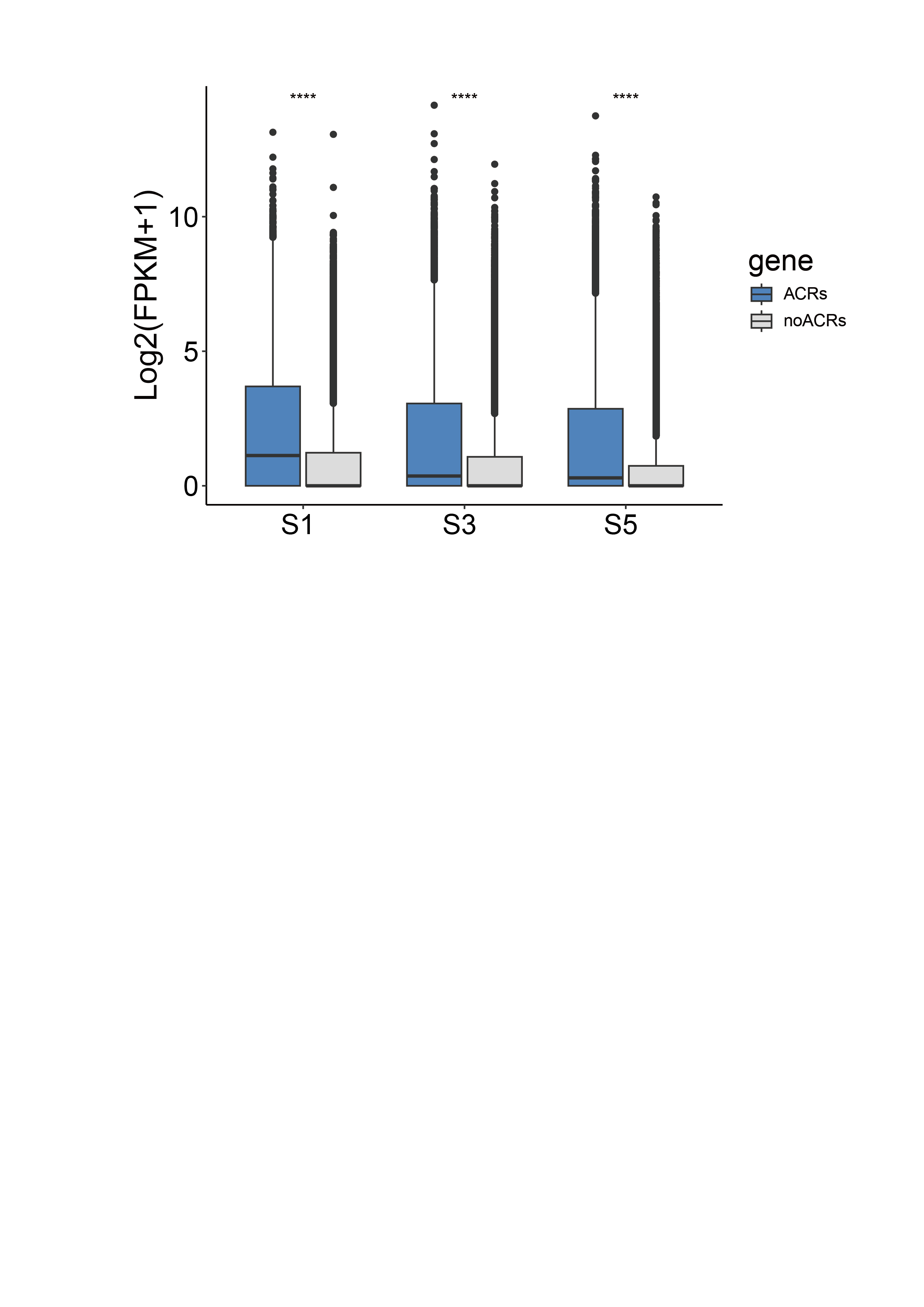

**Fig. S9** Expression levels of genes with or without related accessible regions (ACRs) in root development.

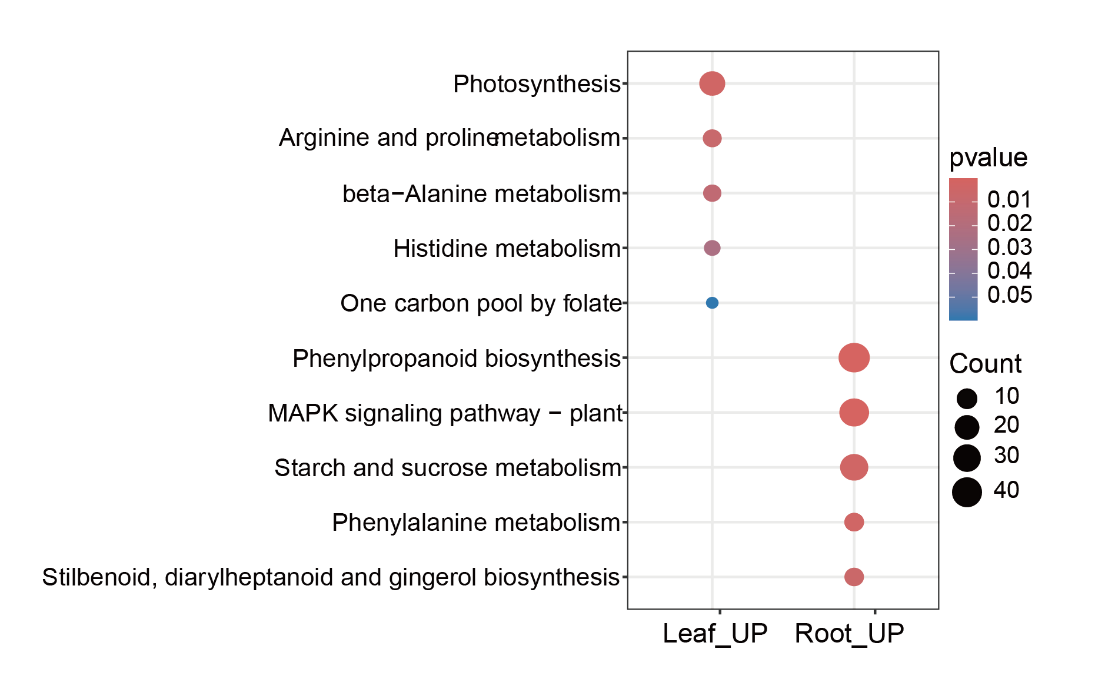

**Fig. S10** KEGG enrichment analysis of differential accessible regions (DARs) up-regulated in leaves (left) and root (right) of *A. dahurica***.
Table S1** Sequences generated in this study.

|  | | | **Base (bp)** | **Depth (×)** |
| --- | --- | --- | --- | --- |
| **DNA** | **PacBio** | | 85,862,285,723 | ~18 |
|  | **Hi-C** | | 328,926,579,300 | ~70 |
| **RNA** | **Illumina** | **Root_rep1** | 52,392,198 | ~11 |
|  |  | **Root_rep2** | 54,644,086 | ~11 |
|  |  | **Root_rep3** | 50,262,761 | ~10 |
|  |  | **Flower_rep1** | 56,513,981 | ~11 |
|  |  | **Flower_rep2** | 59,591,079 | ~12 |
|  |  | **Flower_rep3** | 50,262,761 | ~10 |
|  |  | **Stem_rep1** | 54,299,236 | ~11 |
|  |  | **Stem_rep2** | 58,481,599 | ~12 |
|  |  | **Stem_rep3** | 49,586,093 | ~10 |
|  |  | **Mature _leaf_rep1** | 54,406,096 | ~11 |
|  |  | **Mature _leaf_rep2** | 52,984,058 | ~11 |
|  |  | **Mature _leaf_rep3** | 56,603,962 | ~12 |
|  |  | **Young_leaf_rep1** | 53,074,317 | ~11 |
|  |  | **Young_leaf_rep2** | 50,437,349 | ~10 |
|  |  | **Young_leaf_rep3** | 54,613,159 | ~11 |
|  |  | **S1_rep1** | 48,305,330 | ~10 |
|  |  | **S1_rep2** | 54,633,251 | ~11 |
|  |  | **S1_rep3** | 53,632,578 | ~11 |
|  |  | **S2_rep1** | 43,101,499 | ~9 |
|  |  | **S2_rep2** | 48,832,901 | ~10 |
|  |  | **S2_rep3** | 45,668,244 | ~9 |
|  |  | **S3_rep1** | 49,919,918 | ~10 |
|  |  | **S3_rep2** | 48,329,470 | ~9 |
|  |  | **S3_rep3** | 51,288,449 | ~11 |
|  |  | **S4_rep1** | 49,779,770 | ~10 |
|  |  | **S4_rep2** | 53,141,543 | ~11 |
|  |  | **S4_rep3** | 49,630,061 | ~10 |
|  |  | **S5_rep1** | 49,698,208 | ~10 |
|  |  | **S5_rep2** | 52,724,835 | ~11 |
|  |  | **S5_rep3** | 50,135,167 | ~10 |
|  |  | **S6_rep1** | 52,548,098 | ~11 |
|  |  | **S6_rep2** | 51,440,650 | ~11 |
|  |  | **S6_rep3** | 52,891,402 | ~11 |
| **ATAC** | **Illumina** | **Root** | 39,340,836 | ~9 |
|  |  | **Leaf** | 53,992,304 | ~11 |
|  |  | **S1_rep1** | 33,540,816 | ~7 |
|  |  | **S1_rep2** | 88,318,468 | ~18 |
|  |  | **S3_rep1** | 62,477,708 | ~13 |
|  |  | **S3_rep2** | 61,167,926 | ~13 |
|  |  | **S5_rep1** | 73,942,922 | ~15 |
|  |  | **S5_rep2** | 117,139,522 | ~24 |

**Table S2** Assembly and annotation statistics of the Angelica dahurica genome.

| **Total assembly size (bp)** | 4897920425 |
| --- | --- |
| **Total scaffold number** | 146 |
| **Maximum scaffold length (bp)** | 506786926 |
| **Scaffold N50 (bp)** | 472440809 |
| **Scaffold N90 (bp)** | 353591046 |
| **GC content (%)** | 35.60 |
| **BUSCO# (%)** | 97.20 |
| **LAI** | 20.81 |

#BUSCO anotation using viridiplantae_odb10
**Table S3** The mapping rates of A. dahurica in different tissues.

| **Tissue** | **Read Numbers** | **Mapping Rate** |
| --- | --- | --- |
| S1-1 | 48,305,330 | 95.36% |
| S1-2 | 54,633,251 | 95.02% |
| S1-3 | 53,632,578 | 95.44% |
| S2-1 | 43,101,499 | 94.17% |
| S2-2 | 48,832,901 | 93.91% |
| S2-3 | 45,668,244 | 93.90% |
| S3-1 | 49,919,918 | 95.41% |
| S3-2 | 48,329,470 | 95.49% |
| S3-3 | 51,288,449 | 95.61% |
| S4-1 | 49,779,770 | 95.56% |
| S4-2 | 53,141,543 | 95.62% |
| S4-3 | 49,630,061 | 95.63% |
| S5-1 | 49,698,208 | 95.47% |
| S5-2 | 52,724,835 | 95.15% |
| S5-3 | 50,135,167 | 95.29% |
| S6-1 | 52,548,098 | 94.68% |
| S6-2 | 51,440,650 | 95.23% |
| S6-3 | 52,891,402 | 95.62% |
| R-1 | 52,392,198 | 95.30% |
| R-2 | 54,644,086 | 94.81% |
| R-3 | 50,262,761 | 96.01% |
| S-1 | 54,299,236 | 95.51% |
| S-2 | 58,481,599 | 96.15% |
| S-3 | 49,586,093 | 95.44% |
| ML-1 | 54,406,096 | 83.84% |
| ML-2 | 52,984,058 | 81.51% |
| ML-3 | 56,603,962 | 84.29% |
| YL-1 | 53,074,317 | 86.66% |
| YL-2 | 50,437,349 | 91.48% |
| YL-3 | 54,613,159 | 92.73% |
| F-1 | 56,513,981 | 95.84% |
| F-2 | 59,591,079 | 96.05% |
| F-3 | 50,262,761 | 96.01% |

**Table S4** Statistics of Angelica dahurica pseudomolecules.

| **Chromosome** | **Total length (bp)** | **GC content (%)** | **Gene number** |
| --- | --- | --- | --- |
| Chromosome 1 | 506786926 | 35.67 | 6000 |
| Chromosome 2 | 501082715 | 35.69 | 6447 |
| Chromosome 3 | 490277005 | 35.65 | 5806 |
| Chromosome 4 | 479219616 | 35.61 | 5773 |
| Chromosome 5 | 472440809 | 35.75 | 6726 |
| Chromosome 6 | 432978375 | 35.7 | 5918 |
| Chromosome 7 | 401735355 | 35.75 | 4860 |
| Chromosome 8 | 400149488 | 35.69 | 4730 |
| Chromosome 9 | 390539985 | 35.75 | 5215 |
| Chromosome 10 | 353591046 | 35.69 | 5005 |
| Chromosome 11 | 335116718 | 35.76 | 4586 |
| Unmapped | 134002387 | 35.59 | 353 |
| Total | 4897920425 | 35.60 | 61419 |

**Table S5** Annotation statistics of the Angelica dahurica genome.

| Gene number | 61419 |
| --- | --- |
| Average number of exons per gene | 3.93 |
| Total exon length (Mb) | 63921232 |
| Average exon length (bp) | 264.69 |
| Average number of introns per gene | 2.93 |
| Total intron length (Mb) | 109246157 |
| Average intron length (bp) | 606.68 |
| BUSCO^#^ (%) | 91.00 |
| Number of genes annotated in NR | 53412 |
| Number of genes annotated in KEGG | 31799 |
| Number of genes annotated in Pfam | 35177 |
| Number of genes annotated in GO | 26042 |

#BUSCO anotation using viridiplantae_odb10
**Table S6** Repeat contents in the Angelica dahurica genome.

| **Class** | **Superfamily** | **Count** | **Length (bp)** | **Proportion** |
| --- | --- | --- | --- | --- |
| LTR | Copia | 1216361 | 1417643068 | 28.95% |
| LTR | Gypsy | 911300 | 1187106809 | 24.24% |
| LTR | unknown | 1560595 | 1195529872 | 24.41% |
| TIR | CACTA | 232178 | 97986548 | 2.50% |
| TIR | Mutator | 271674 | 104900806 | 1.40% |
| TIR | PIF_Harbinger | 59310 | 16859532 | 0.29% |
| TIR | Tc1_Mariner | 208479 | 63621194 | 0.82% |
| TIR | hAT | 168495 | 56472214 | 0.99% |
| nonTIR | helitron | 67893 | 23853899 | 3.47% |
| Total |  | 4966673 | 4264156825 | 87.08% |

**Methods S1** Genome size estimation.

Both k-mer and flow cytometry methods were used for *A. dahurica* genome size estimation. Fresh leaves were vertically chopped to release nuclei in cold LB01 lysis buffer (15 mmol/L Tris, 2 mmol/L Na2EDTA, 0.5 mmol/L spermine tetrahydrochloride, 80 mmol/L KCl, 20 mmol/L NaCl, 0.1% (v/v) TritonX-100, 15 mmol/L β-mercaptoethano, pH 7.0~8.0), with a disposable razor blade. The nuclei suspension was filtered through a 40 μm cell strainer, stained with 20 μg mL-1 propidium iodide (DNA fluorochrome; Thermo Fisher Scientific, Waltham, MA, USA) and 20 μg mL-1 RNase A (Thermo Fisher Scientific), and ice-bathed for 30 min in the dark. The fluorescence intensity of stained nuclei was analyzed with CytoFLEX (Beckman Coulter, Miami, FL, USA). The value of nuclear DNA was calculated by comparing the nuclear peaks on a linear scale with the peak for *Foeniculum vulgare* Mill. (1C=1.01G) and *Angelica sinensis* (1C=2.37G) using CytExpert software v.2.3 (Beckman Coulter, Indianapolis, IN, USA). Clean whole genome sequence (WGS) reads were analyzed to obtain the 19-mer distribution with Jellyfish v2.3.0 (Marcais & Kingsford, 2011). The output file was used as the input for GenomeScope 1.0 (Vurture *et al.*, 2017) to estimate the genome size and heterozygosity rate.

**Methods S2** DNA and RNA preparation and sequencing.

Fresh leaves of *Angelica dahurica* were harvested for the extraction of high molecular weight (HMW) genomic DNA with the DNeasy Plant Mini Kit (Qiagen, USA). A quantity exceeding 50 µg of HMW DNA was used for the construction of SMRTbell^TM^ libraries. These libraries were sequenced on the PacBio Sequel II platform with the circular consensus sequencing (CCS) mode. For the Hi-C approach, plant tissues underwent formaldehyde treatment for fixation, with the cross-linked DNA subsequently digested by DpnII throughout the night. The digested fragments' sticky ends were then biotinylated and ligated in a random fashion. These chimeric fragments, reflecting the original cross-linked physical interactions, were further processed into paired-end sequencing libraries. The sequencing phase for these libraries was conducted on the Illumina NovaSeq 6000 platform, generating 2 x 150 bp reads.

Concurrently, total RNA was extracted from the roots across seven different growth periods, in addition to flowers, young leaves, old leaves, and stems, employing the RNAprep Pure Plant Kit (TIANGEN, Beijing, China). cDNA synthesis was carried out with 20 µg total RNA, Rever Tra Ace (TOYOBO, Osaka, Japan) and oligo (dT) primers, adhering to the instructions of the user manual. Subsequently, RNA-Seq reads were obtained on the Illumina NovaSeq 6000 platform.

**Methods S3** Genome assembly and annotation.

The genome assembly of *A. dahurica* was achieved by combining data from PacBio CCS and Hi-C technologies, employing Hifiasm v0.15.5-r350 software with default parameters (Table S1) (Cheng *et al.*, 2021). Subsequently, Purge Haplotigs v1.1.1 (Roach *et al.*, 2018) was used to remove redundant sequences because of a very high heterozygosity observed in *A. dahurica* genome. Next Quality-controlled Hi-C reads were then aligned to the contig assembly of *A. dahurica* with Juicer (Durand *et al.*, 2016). A preliminary chromosome-level assembly was generated automatically with the 3D-DNA v180114 pipeline to rectify wrong assemblies, orders and orientations (Dudchenko *et al.*, 2017). Further refinement and validation of this assembly were conducted manually through the Juicebox Assembly Tools (https://github.com/aidenlab/Juicebox), a crucial step in enhancing the accuracy of assembly. Finally, the quality of the chromosome-level genome assembly was rigorously assessed with Benchmarking Universal Single-Copy Orthologs (BUSCO v5.1.2) and LTR Assembly Index (LAI) (Ou *et al.*, 2018; Seppey *et al.*, 2019).

We employed the EDTA v2.0.1 (Ou *et al.*, 2019) for the comprehensive identification of transposable elements, encompassing LTR, TIR, and non-TIR elements. To predict coding gene structures within the repeat-masked genome, we utilized a multifaceted approach that included *ab initio* predictions, evidence from homologous proteins, and transcriptome data. For the *de novo* prediction of protein-coding genes, we used AUGUSTUS v.2.3.3 (Stanke *et al.*, 2006). MAKER v3.01.03 pipeline (Cantarel *et al.*, 2008) was used to annotate gene structures with RNA-seq data from 11 tissues, including roots from seven developmental stages, flower, young leaves, old leaves, stems, in our study and protein sequences from species of *Daucus carota* and *Ligusticum chuanxiong*. Further annotations of protein-coding genes were conducted by GFAP (Xu *et al.*, 2023) and EGGNOG-MAPPER v.1.0.3 (Cantalapiedra *et al.*, 2021) to the Kyoto Encyclopedia of Genes and Genomes (KEGG, https://www.kegg.jp), Gene Ontology (GO, http://geneontology.org) and PFAM (http://pfam.xfam.org/) databases.

**Methods S4** Phylogenetic analyses.

Paralogs and orthologs were identified among seven plant species: *Centella asiatica* (L.) Urban (Pootakham *et al.*, 2021), *Bupleurum chinense* DC (Zhang *et al.*, 2022), Daucus carota L. (Iorizzo *et al.*, 2016), *A. dahurica*, *A. sinensis* (Han *et al.*, 2022) and *Ligusticum chuanxiong* haplotype A (Nie *et al.*, 2024) with the OrthoFinder v2.5.2 (Emms & Kelly, 2019), and protein sequences of single-copy orthologous genes were used to construct a phylogenetic tree. The concatenated amino acid sequences were aligned using MAFFT v7.271 (Katoh & Standley, 2013) and trimmed with trimAI v1.4.rev22 (Capella-Gutierrez *et al.*, 2009). A maximum likelihood phylogenetic tree was constructed using RAxML v.8.2.12 of a PROTGAMMAJTT model with 1000 bootstrap replicates (Stamatakis, 2014), and *C. asiatica* was used as the outgroup. The species tree was then used as an input to estimate divergence time in the MCMCTree program of the PAML package (Yang, 2007). Fossil time of divergence between B. chinense and D. carota was used for time calibrations from TIMETREE (<http://www.timetree.org/>). The expansion and contraction of gene families were inferred with CAFE5 (Mendes *et al.*, 2020) based on the chronogram of the above-mentioned nine plant species.

Syntenic analyses and gene duplication identification Syntenic blocks within one species or between two species were defined by MCscanX (Wang *et al.*, 2012) based on homologous gene sets using BLASTP v2.10.0 (E-value < 1e-5; the number of genes required to call a syntenic block ≥ 5). To further identify the pattern of genome-wide duplications in *A. dahurica*, duplicated genes were divided into five categories: whole-genome duplication (WGD), tandem duplication (TD), proximal duplication (PD), transposed duplication (TRD), dispersed duplication (DSD), using DupGen_Finder v1.12 (Qiao *et al.*, 2019) with the default parameters. Genes in the five duplicate categories were further fed with Kyoto Encyclopedia of Genes and Genomes (KEGG) terms enrichment analysis with TBtools v2.030 (Chen *et al.*, 2023).

**Methods S5** Multi-omics mining for candidate CYP450 genes.

To identify potential CYP450 genes in A. dahurica, we employed two distinct approaches. 1) We conducted a search based on conserved domains using HMMER v3.3 (Finn *et al.*, 2015); 2) we utilized a sequence similarity approach, employing the protein sequence of the CYP450 enzyme CYP71AZ4, previously characterized in Pastinaca sativa (Apiaceae), as a query sequence for local BLAST in the A. dahurica genome (--evalue 1e-6). After the exclusion of potential pseudogenes (predicted protein sequences < 300 amino acids), we considered the intersecting sequences obtained from both methods as putative CYP450 genes in A. dahurica. Next, we utilized a comprehensive approach that incorporated gene expression analysis, metabolite content assessment, and phylogenetic relationships to screen two candidate genes for P8H and four candidate genes for P5H.

To narrow down the set of candidate genes, a phylogenetic analysis of these sequences with CYP450 protein sequences in *Arabidopsis thaliana*, as well as sequences verified to be involved in furanocoumarin synthesis. All sequences were aligned by MAFFT v7.487 (Katoh *et al.*, 2019). Phylogenetic reconstructions were implemented in IQtree v2.2.5 (Minh *et al.*, 2020) to infer the maximum-likelihood (ML) tree. The final tree was visualized and annotated in iTOL v6.7.6 (https://itol.embl.de/) (Letunic & Bork, 2021). With referrence to CYP450 sequences of *A. thaliana*, we categorized the phylogenetic tree into seven clades: CYP71 Clan, CYP86 Clan, CYP97 Clan, CYP72 Clan, CYP711 Clan, CYP85 Clan, and CYP51 Clan. Based on the phylogenetic relationships, we initially screened the 25 *A. dahurica* genes located in CYP71 Clan and clustered with CYP71AZ were identified as preliminary candidates for P8H and P5H (Fig.S6). We further annotated these 25 preliminary candidates with matrices of expression levels from multiple tissues and from different root developmental periods of *A. dahurica*. Ultimately, two genes with Pearson correlation coefficients greater than 0.75 with xanthotoxol content and four genes with Pearson correlation coefficients greater than 0.75 with bergaptol content were considered as candidate genes for experimental validation.
